# FoxP3 forms a head-to-head dimer in vivo and stabilizes its multimerization on adjacent microsatellites

**DOI:** 10.1101/2025.09.16.676304

**Authors:** Fangwei Leng, Ryan Clark, Wenxiang Zhang, Thibault Viennet, Haribabu Arthanari, Wang Xi, Sun Hur

**Author notes:** These authors contributed equally to this work (Fangwei Leng, Ryan Clark).

## Abstract

FoxP3, the master regulator of Tregs, employs two DNA-binding modes to recognize diverse DNA sequences. It multimerizes on long TnG repeats (n = 2–5) to bridge DNA segments and stabilize chromatin loops, and it forms head-to-head (H-H) dimers on inverted repeat forkhead motifs (IR-FKHM) without bridging DNA. Although genomic data confirm its multimeric role, in vivo evidence for H-H dimerization has been elusive. Here, unbiased pull-down sequencing uncovers a range of relaxed motifs that drive H-H dimerization, enabling systematic genome-wide analysis. We demonstrate that FoxP3 binds genomic DNA as both H-H dimers and multimers in Tregs, with H-H binding often seeding and stabilizing multimerization on adjacent TnG repeats—especially on shorter, suboptimal repeats. While multimerization is conserved across FoxP family members, H-H dimerization is unique to FoxP3 orthologs, conferred by its divergent accessory loop. This dual-mode strategy broadens FoxP3’s sequence repertoire and enhances its architectural function in chromatin looping.

## Introduction

Transcription factors (TFs) are sequence-specific DNA-binding proteins central to virtually all gene regulatory networks. Of the ∼1,600 TFs characterized to date, about half contain a single DNA-binding domain^1,2^ and have long been thought to adopt a single, conserved DNA-binding mode. However, emerging evidence reveals a more complex landscape. One example is FoxP3, which was recently shown to adopt multiple conformations depending on the DNA sequence^3–5^. The relationships and functional distinctions between these distinct binding modes, however, remain unclear.

FoxP3 belongs to the forkhead family, whose members adopt the conformation of a winged-helix forkhead domain (FKH) to bind DNA^6^. Although early reports suggested otherwise^7,8^, FoxP3’s forkhead domain also assumes this canonical fold^3^. Unlike most forkhead TFs, however, FoxP3 binds poorly to isolated consensus forkhead motifs (FKHMs)^3–5^. Instead, it requires composite recognition sites—either an inverted repeat with a 4-nt gap (IR-FKHM)^3^ or TnG repeat microsatellites^4,5,9^. In both cases, a loop adjacent to FKH, namely Runx1-binding region (RBR, Figure 1A), plays a critical role as it mediates FoxP3-FoxP3 interactions^3,10^. On IR-FKHM, FoxP3 forms a head-to-head (H-H) dimer through the RBR-RBR interactions^3^. On tandem repeats of TnG sequence (n = 2-5), FoxP3 forms complex multimers in various stoichiometries and configurations, using both RBR and FKH^4,5^. The multimerization bridges 2-4 DNA segments^4,5^ and stabilizes chromatin loops in vivo^4,11,12^. Although FoxP3 is constitutively dimeric through its coiled coil (CC) domain^13^ (Figure 1A), these additional RBR-mediated interactions are essential for cooperative binding to IR-FKHM and TnG repeats^3–5^.

**Figure 1.**
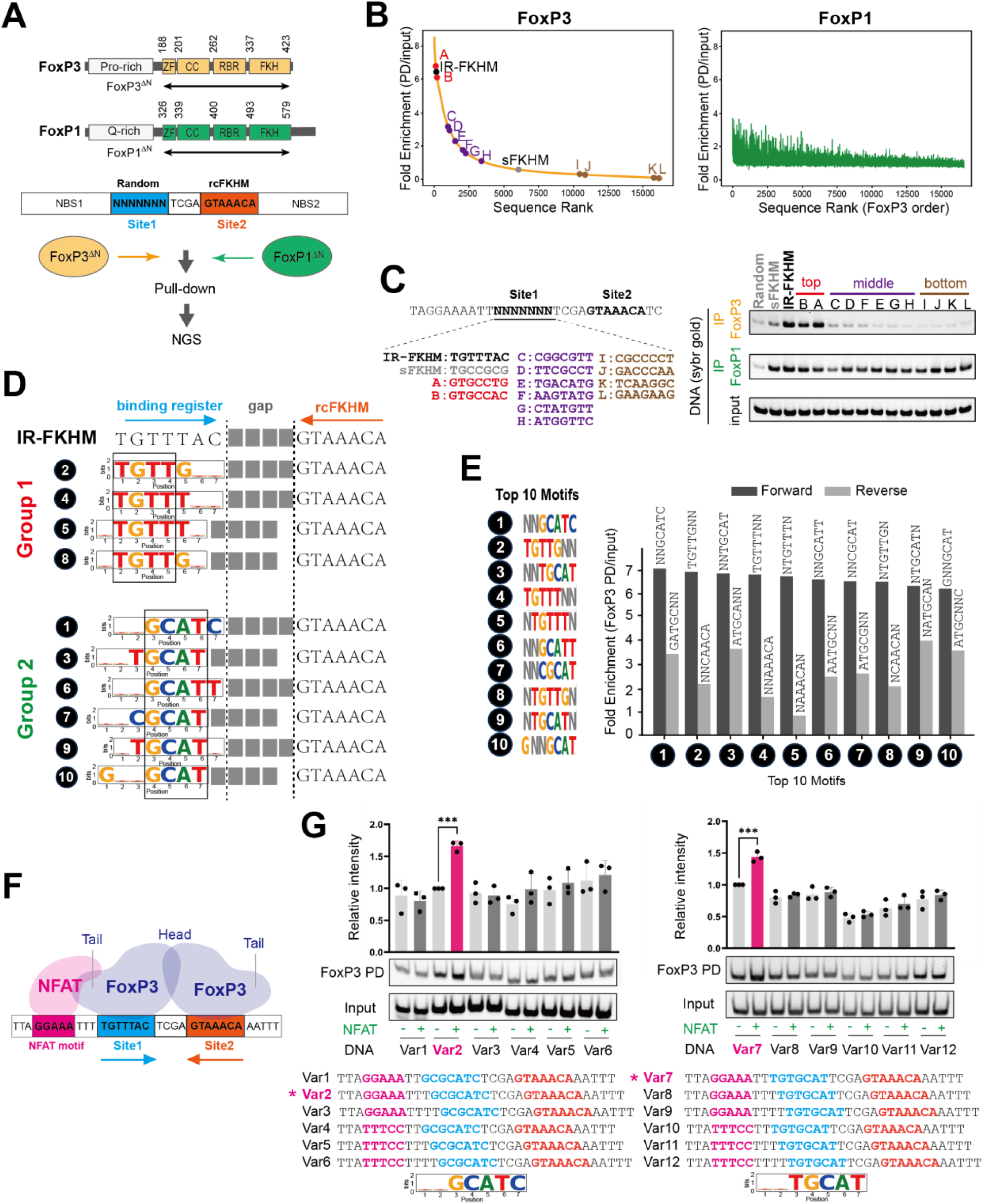
FoxP3 recognizes diverse DNA motifs paired with FKHM through H-H dimerization. A. Top: domain architecture of FoxP3 and FoxP1 (CC, coiled-coil domain; ZF, zinc finger domain; RBR, Runx1 binding region). Bottom: schematic of pull-down (PD)-seq. Degenerate DNA library (80 bp) used for PD-seq harbors a 7-nt random sequence (site 1) followed by a 4-nt gap and a 7-nt reverse complement FKHM (rcFKHM, site 2), flanked by non-binding sequences (NBS1 and NBS2, see Table S1 for sequence). MBP-tagged, N-terminal deletion constructs (FoxP3^ΔN^ and FoxP1^ΔN^) were used for pull-down. Co-purified DNA was analyzed by next-generation sequencing (NGS). Same experimental condition was used for FoxP3 and FoxP1. B. Fold enrichment (PD/input) for each sequence from the random-rcFKHM oligo PD-seq in (A). Both FoxP3 PD (left) and FoxP1 PD (right) samples were ranked in the descending order of FoxP3 fold enrichment. C. Selected sequences from the top (red), middle (purple), and bottom (brown) ranked groups in (B) were further tested by FoxP3/FoxP1 pull-down, followed by analysis of the co-purified DNA on native gel (SybrGold stain). IR-FKHM served as a positive control. Random, non- binding sequence and sFKHM—comprising a non-binding sequence in site 1 paired with rcFKHM in site 2—served as negative controls for FoxP3 binding throughout this study. See Table S1 for sequences. Same experimental condition was used for FoxP3 and FoxP1. D. Top 10 motifs from FoxP3 PD-seq allowing degeneracy at two bases (see Figure S1E for justification). These motifs fall into two classes: group 1 resembling FKHM (TGTTT or TGTTG), and group 2 characterized by a core sequence of GCAT. Numbers next to individual motifs indicate rank orders based on fold enrichment in (E). E. FoxP3 PD-seq fold enrichment for sequences harboring the top 10 motifs in site 1 in forward (black) or reverse (grey) orientation. F. NFAT–FoxP3 cooperative binding occurs when the NFAT site is positioned 3 nt upstream of FKHM in site 1, allowing direct interaction between NFAT protein and FoxP3 tail. G. NFAT–FoxP3 cooperativity analysis. DNAs containing group 2 motifs with varying NFAT site positions and orientations (0.1 μM) were incubated with MBP-tagged FoxP3^ΔN^ (0.4 μM) with and without NFAT (0.2 μM), followed by MBP pull-down and gel analysis of co-IPed DNA (SybrGold stain). NFAT–FoxP3 cooperativity was assessed by the enhancement of FoxP3– DNA binding upon addition of NFAT. Data are representative of three biological replicates and presented as mean ± s.d. p-values were calculated for unpaired two-tailed t-test; *** p < 0.001.

A central question is whether these in vitro-observed conformations occur in vivo. Genomic analyses of FoxP3-occupied loci––via CNR-seq^12,14^, ChIP-seq^14,15^ or ChIP-exo^9^––have consistently showed strong enrichment of TnG repeats^4,5,9^, supporting a model where FoxP3 multimerizes to stabilize chromatin loops in Tregs. By contrast, IR-FKHM motifs associated with H-H dimerization were previously reported to show only minimal enrichment within a relatively limited but highly reproducible set of FoxP3-bound regions^3^. Functionally disentangling H-H dimerization from multimerization using mutagenesis has also provden difficult. This is because their interaction interfaces overlap—while some residues involved in multimerization lie outside the RBR, those critical for H-H dimerization, which are mainly within the RBR, are also essential for multimerization^4,5^. Notably, while multimerization on TnG is common to all four members of FoxP family (FoxP1-4)^4^, H-H dimerization is unique to FoxP3^3^, raising the question of whether it reflects an evolutionarily adopted function specific to FoxP3.

We hypothesized that H-H dimerization occurs in vivo, but was undetected because it takes place on DNA sequences beyond IR-FKHM, and/or because previous analyses of in vivo occupancy data were too restrictive. To test these possibilities, we conducted a more systematic in vitro pull-down sequencing together with broader genomic data analyses. These new results revealed that FoxP3 recognizes a wider range of motifs as a H-H dimer than previously appreciated, and that these H-H motifs are enriched within FoxP3 genomic targets in vivo. These H-H sites often lie immediately adjacent to TnG repeats, where H-H dimerization assists FoxP3 multimerization on short, suboptimal TnG repeats. Together, these findings uncover a previously unrecognized role for H-H dimerization in expanding FoxP3’s architectural functions and provide new mechanistic insight into its mode of action.

### 1. FoxP3 recognizes diverse DNA motifs paired with FKHM through H-H dimerization

To capture the full repertoire of FoxP3’s sequence preferences, we performed an unbiased pull-down assay followed by deep sequencing (PD-seq) using a library of degenerate DNA oligonucleotides. Unlike our previous genome-based PD-seq^4^—which predominantly identified TnG repeats due to their genomic abundance^16^ and the large fragment sizes that favor multimerization^4^—we here specifically aimed to isolate the dimeric binding mode. We designed oligos containing a randomized 7-nt segment (site 1) followed by a 4-nt spacer and a fixed reverse-complement FKHM (rcFKHM, site 2), all flanked by 27 nt upstream (NBS1) and 35 nt downstream (NBS2) of non-binding sequence (Figure 1A; Table S1). Note that if site 1 were an FKHM, this design would recreate an IR-FKHM that facilitates FoxP3 H-H dimerization^3^. For clarity, all sequences are reported on the forward strand.

Recombinant FoxP3 protein fused with an maltose-binding protein (MBP) tag was used to pull down via amylose resin as in the previous study^4^. Since full-length FoxP3 could not be purified due to solubility issue and non-specific proteolytic cleavage, we used a construct lacking the N-terminal disordered region (residue 188-423, FoxP3^ΔN^). Both the MBP tag and the truncation of the N-terminal portion has little impact on DNA sequence specificity^3,4^. MBP-FoxP1^ΔN^ (residue 326-579) construct with the equivalent boundary was used for comparison (Figure 1A).

We first ranked individual sequences based on read counts normalized to input (Figure 1B, see the full sequence list in Table S2). The two biological replicates of FoxP3-PD-seq were highly concordant (Figure S1A), whereas FoxP3 and FoxP1 PD-seq displayed a striking divergence (Figure 1B, S1B). While FoxP3 binding strongly depended on site 1 sequence, FoxP1 exhibited markedly lower reliance on this region. Testing selected sequences from the top-, middle-, and bottom-ranked groups based on the FoxP3 PD-seq further confirmed this conclusion (Figure 1C). These findings suggest that FoxP3 recognizes both site 1 and site 2, whereas FoxP1 engages only the rcFKHM in site 2. Moreover, minimal enrichment of FKHM in site 1 for FoxP1––despite its known ability to bind isolated FKHM––implies that site 1 occupancy is either incompatible with site 2 co-occupancy or not needed for FoxP1.

To identify motif preference, we initially employed conventional de novo motif discovery tools, such as MEME^17^. However, whether applied to the entire 18 nt region encompassing site 1, a 4- nt gap and rcFKHM or just site 1, MEME identified few discernible sequence features (Figures S1C, S1D). Instead, we systematically analyzed reads per million (RPMs) for each unique sequence of site 1. Allowing degeneracy at 2 out of 7 bases (2N motifs) markedly simplified the representation of highly enriched sequences, enabling 10 motifs to collectively account for the sequences with top 10% of enrichment values (Figure S1E, see Table S3 for the top 25 motifs). Pulldown assays of representative sequences confirmed that all 10 motifs are comparable to IR-FKHM in FoxP3 binding (Figure S1F). These 10 motifs were classified into two groups: (1) those containing TGTTT or TGTTG—closely resembling FKHM (TGTTTAC)—and (2) those containing GCAT flanked by either T or C on each side, akin to a non-canonical FoxM1 motif^18^ but previously unrecognized for FoxP3 (Figure 1D).

Notably, all top 10 motifs paired with FKHM displayed a strong orientational bias (Figure 1E), suggesting that FoxP3’s binding to site 1 depends on its binding to site 2 sequence rcFKHM. We hypothesized that this bias reflects FoxP3’s H-H dimerization. Supporting this hypothesis, group 1 (G1) sequences in site 1 align with FKHM, forming sequences similar to IR-FKHM with a gap of 3-4 nt (Figure 1D). That is, motifs #2 and #4 in site 1 recreate IR-FKHM with a 4-nt gap, whereas motifs #5 and #8 generate 3-nt gapped variants (Figure 1D). This is consistent with previous observations that FoxP3 prefers IR-FKHM with a 4 nt-gap, but can tolerate a 3-nt gap^3^.

To test whether FoxP3 also binds G2-rcFKHM in an H-H orientation, we employed a FoxP3-NFAT cooperativity assay^3^, which exploits NFAT’s ability to dock onto FoxP3’s “tail” only when their DNA sites are correctly phased^7,8^ (Figure 1F). In this assay, a 7-nt test sequence placed 3 nt downstream of the NFAT site (GGAAA) will restore cooperativity if FoxP3 engages that sequence in the same register as FKHM. When we tested GC<u>GCAT</u>C, we observed NFAT-dependent enhancement of FoxP3–DNA binding exclusively at a 3-nt offset—and only in the forward orientation (Figure 1G, left). When the core GCAT was shifted by 1 nt in another G2 sequence TGT<u>GCAT,</u> optimal cooperativity was observed at a 2-nt offset, again only in the forward orientation (Figure 1G, right). These results suggest that FoxP3 binds NN<u>GCAT</u>N motif in the same manner and register as FKHM, and the enriched sequences are consistent with H-H dimerization with a 4-nt gap.

Altogether, these findings suggest that FoxP3 forms a H-H dimer on a broader set of DNA sequences, including TGTTNNN or NNGCATN paired with rcFKHM at a 4 or 3 nt-gap, and that this property is specific to FoxP3 and not shared with FoxP1.

### 2. FoxP3 recognizes pairs of group 1 and 2 sequences in H-H configuration

Next, we examined how FoxP3 binds DNA when both sites 1 and 2 were allowed to vary, rather than fixing site 2 to rcFKHM. Specifically, we asked whether FoxP3 could bind the 10 motifs in Figure 1 when paired with each other (without rcFKHM) and whether the orientational bias for the H-H configuration is preserved. To test this, we performed FoxP3 PD-seq using a DNA oligo library based on the construct in Figure 1A, but with degenerate sequences in both sites 1 and 2 (random-random, Figure 2A). Because this expanded library contained 4^14^ (268,435,456) possible sequences—too large for comprehensive input sequencing—we sequenced only the PD-enriched samples to a depth of ∼280 million reads each. For FoxP3, sharp enrichment was seen for fewer than 10,000 unique sequences, which differed vastly from that of FoxP1 PD-seq (Figure 2B), consistent with their difference in DNA specificity.

**Figure 2.**
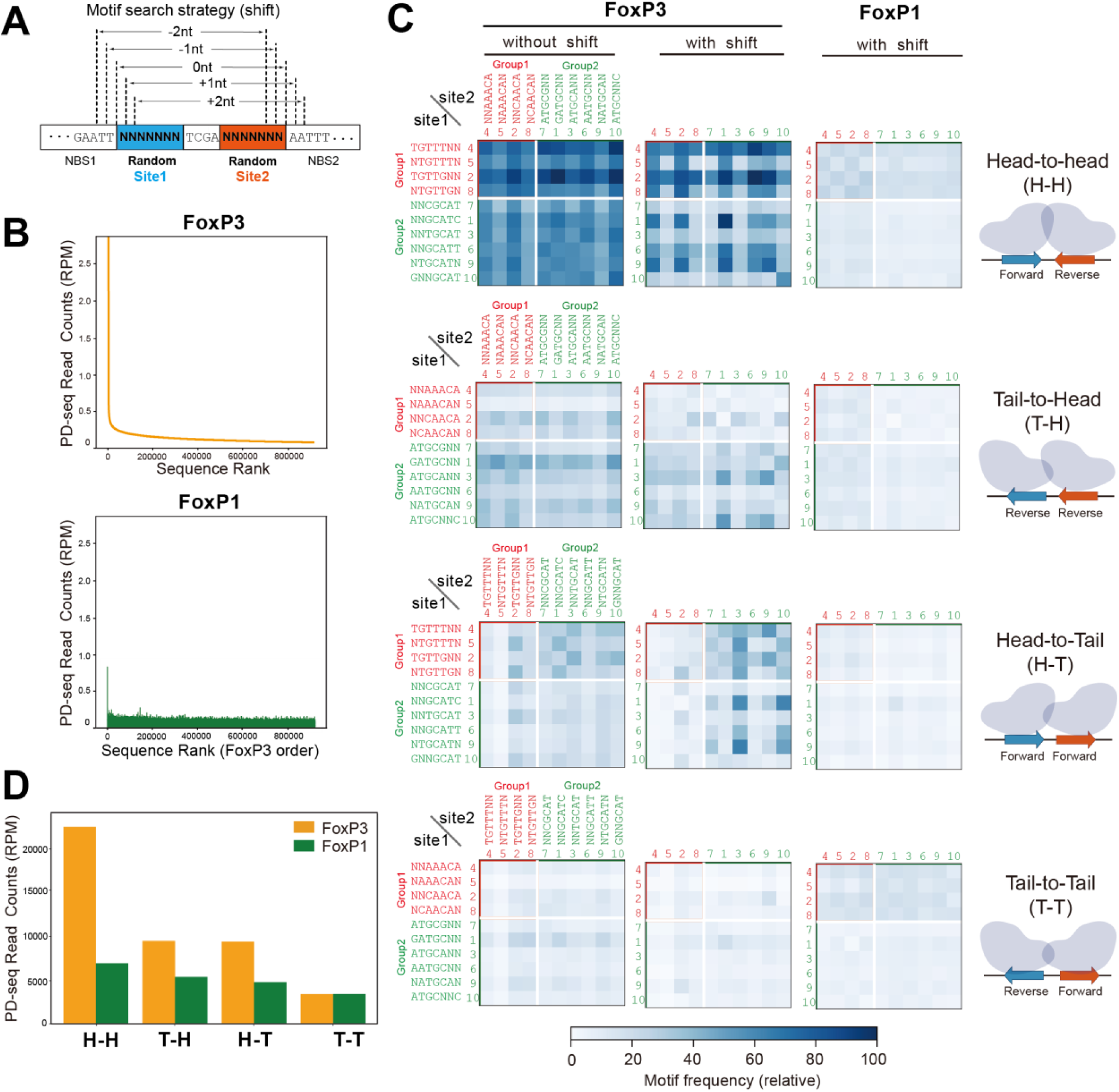
FoxP3 recognizes pairs of group 1 and group 2 sequences in H-H orientation. A. Schematic of the random–random DNA library (80 bp) used for FoxP3/FoxP1 PD-seq. PD-seq was performed as in Figure 1A, except that the library contained degenerate 7-nt sequences in both sites 1 and 2, flanking a 4-nt gap. Since FoxP3 registers were not fixed as for the random–rcFKHM library, motif searches were performed with and without relative positional offsets of −2 to +2 nucleotides between the variable sites. B. FoxP3 (top) and FoxP1 (bottom) PD-seq read counts (in read-per-million, RPM) for sequences in the random-random DNA library. Shown are the top 1% unique sequences, arranged in descending order based on FoxP3 PD-seq read counts. C. Heatmaps showing the frequency of motif pairs in PD-seq. Top 10 motifs––group 1 (red) and group 2 (green)––can be paired in four orientations: head-to-head (H–H), tail-to-head (T–H), head-to-tail (H–T), and tail-to-tail (T–T). Motif pair frequency was quantified by counting sequences in site 1 and site 2 in PD-seq, either without positional shifts (left), or allowing shifts of −2 to +2 nt for FoxP3 (center) or FoxP1 (right). Motif pair frequencies were scaled from 0 to 100 for visualization. For the shifted conditions, frequencies were additionally corrected for theoretical input bias (see Methods). A consistent scale was used for both FoxP3 and FoxP1 after shifting. Numbers next to individual motifs indicate the enrichment rank order from the random-rcFKHM PD-seq in Figure 1E. D. Total paired motif read counts for H-H, T-H, H-T and T-T. All motif counts were from (C) with shifts.

To examine the orientational bias of FoxP3, we generated heatmaps showing relative read counts of each pair of the 10 motifs in sites 1 and 2 in one of the four orientations (H-H, H-T, T- H or T-T). The comparison showed that H-H pairing is indeed preferred, compared to the other configurations (Figure 2C).

Since the binding positions of FoxP3 in the random-random library were no longer fixed (unlike in the random-rcFKHM library), we also allowed for -2, -1, 0, +1, and +2 bp shifts when scanning for motifs (Figure 2A). This, however, inevitably introduces the influence of both flanking and gap sequences. To correct this, we computed the baseline probability of observing each motif pair under random shifts and normalized the observed motif frequency in the PD samples by these baseline probabilities (see Methods). Even after this positional relaxation, H-H dimerization remained markedly favored over other configurations (Figure 2C, 2D). This bias for H-H orientation was more pronounced in FoxP3 than in FoxP1 (Figure 2C, 2D), in agreement with the PD-seq result using the random-rcFKHM oligonucleotides.

Note that random-random PD-seq results were not used for de novo motif discovery due to the ambiguity in FoxP3 register, which can artificially influence motif enrichment. For example, although inverted repeat of GCGYGCH (Y indicates T/C, H indicates A/C/T) appeared enriched in the top 25 sequences from this library (Table S4), this does not represent a bona fide FoxP3 motif. GCGYGCH (or its reverse complement) was not enriched in the random-rcFKHM library (green lines in Figure S1E). Moreover, mutating the spacer or flanking sequences in IR- GCGTGCA abolished FoxP3 binding, whereas similar mutations had minimal impact on IR-FKHM binding (Figure S1G). This suggests that the apparent enrichment of IR-GCGYGCH reflects FoxP3 recognition of flanking or spacer elements rather than the core motif itself. Consistent with this, closer inspection revealed that IR-GCGYGCH within our random-random construct contains two inverted pairs of high-ranking motifs from Figure 1 in ±2 shifted positions (Figure S1H), likely explaining its strong enrichment. Given these complexities and the high risk of over- interpretation, we relied primarily on the random-rcFKHM library for de novo motif discovery, while using the random-random library mainly to validate pairing orientations.

Altogether, these data confirmed H-H dimerization of FoxP3 using pairs of both G1 and G2 sequences and the lack of such dimerization for FoxP1.

### 3. FoxP3 recognizes both H-H motifs and TnG repeats in Tregs

Using the newly identified FoxP3 motifs, we systematically assessed whether H-H motifs are enriched within FoxP3-occupied genomic regions in vivo. We first compared the frequency of individual motifs in FoxP3-bound regions—25,724 peaks defined using four individual FoxP3 Treg ChIP-seq data from two groups^14,15^—with their frequency in open chromatin regions (OCRs) from Treg ATAC-seq^19^ (Figure 3A). This definition of FoxP3-bound regions is intentionally more relaxed than in the previous study^3^, as discussed later. For FoxP3-bound regions, we also examined CUT&RUN-seq (CNR-seq, 18,552 peaks) datasets from Tregs^12,14^ and recent ChIP-exo data^9^ (4,307 peaks), which provide precise FoxP3 footprints, though derived from ectopically expressed FoxP3 in mESCs (Figure S2A). For significantly enriched motif pairs (p < 0.05; black circles in Figure 3A), fold enrichment values were visualized in heatmaps (Figure 3B, Figure S2A). Motif frequencies were calculated only on the positive reference strand; analyzing the negative strand would convert T-H into H-T and invert the rows/columns for H-H and T-T, explaining the similarity between H-T and T-H maps and the diagonal symmetry observed in H-H and T-T maps.

**Figure 3.**
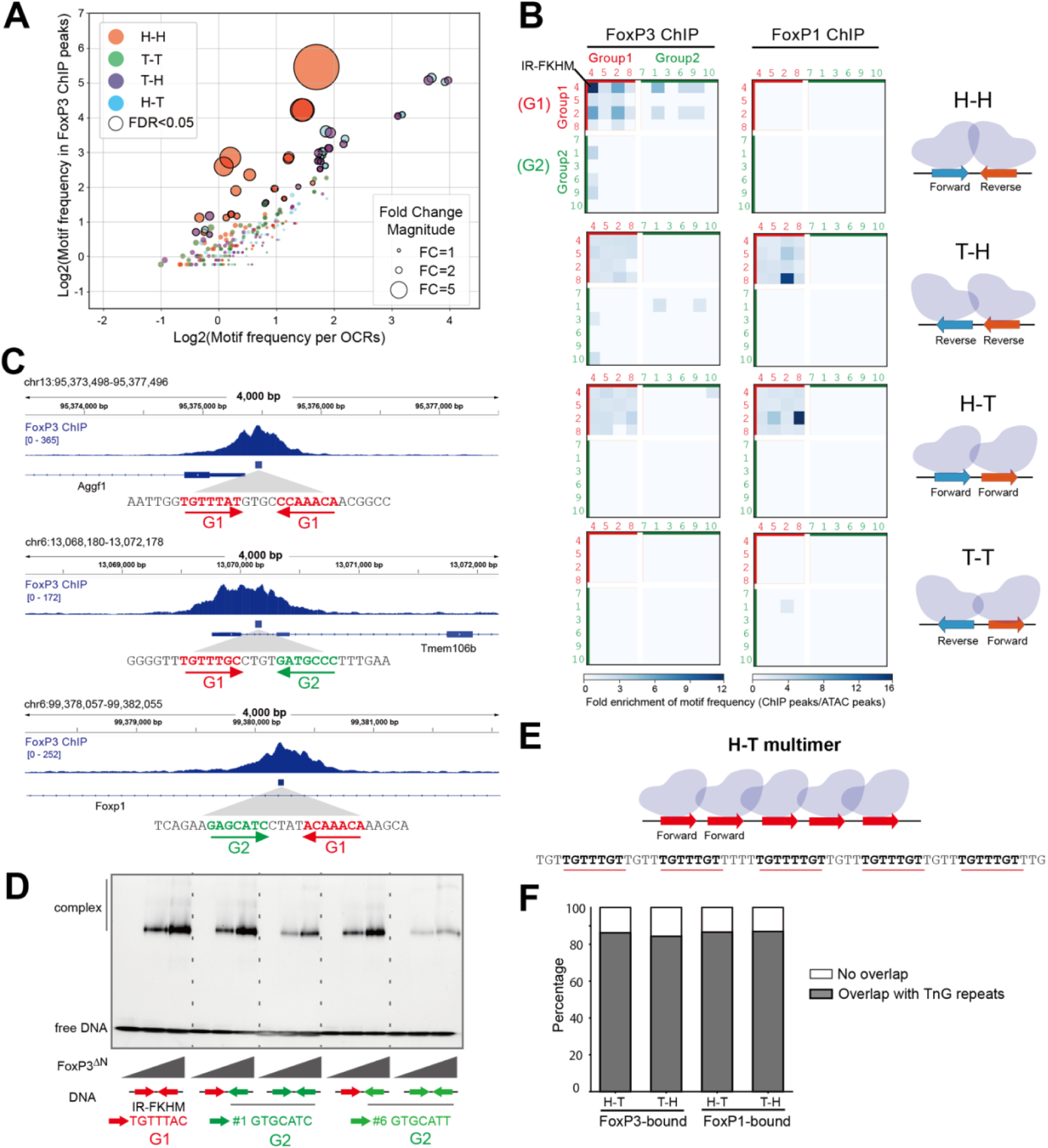
FoxP3 recognizes both H-H motifs and TnG repeats in Tregs. A. Comparison of the log2-transformed motif frequency in FoxP3 ChIP-seq peaks (25,724 peaks) vs. open chromatin regions (OCRs based on ATAC-seq, 502,437 peaks). Each point represents one of the 400 motif pairs (10 motifs x 10 motifs x 4 orientation combinations) with a 4-nt gap, colored by the motif orientation relative to the reference genome (positive strand). Motif frequency was defined as the number of motif occurrence within all ChIP- peaks (or ATAC-peaks) divided by the total length (in million bp) of those peaks. Dot size reflects fold enrichment of motif frequency in ChIP peaks over OCRs. Motif pairs significantly enriched in FoxP3 ChIP peaks (p<0.05, exact binomial test) are outlined in black and are further analyzed in (B). B. **Heatmaps showing fold change of the motif frequency in FoxP3 ChIP-peaks over OCRs, grouped based on the motif pair orientation. The top 10 motifs are further categorized into group 1 (G1, red) and group 2 (G2, green) sequences. FoxP3** ChIP-seq data are from Tregs (left), whereas FoxP1 data are from CD8+ T cells (right). Only the motif pairs p<0.05 are shaded. C. Genome browser views of FoxP3 ChIP-seq peaks harboring different H-H pairs: (i) G1-G1, (ii) G1-G2, and (iii) G2-G1, as indicated in the FoxP3 heatmap in (B). Tracks (top to bottom): FoxP3 ChIP-seq, H-H pair regions within Foxp3 ChIP peaks, and Refseq gene annotation. D. Native gel shift assay of MBP-tagged FoxP3^ΔN^ (0, 0.4, 0.8 μM) with DNAs (0.2 μM) containing different H-H pairs. E. Schematic of a representative TnG repeat region within a FoxP3-bound locus, where TnG repeat was counted as multiple H-T (G1–G1) motif pairs with a 4-nt gap. This configuration facilitates FoxP3 multimerization and DNA bridging. F. Stacked bar plot showing the proportion of enriched H-T/T-H (G1–G1) motif pairs that overlap with previously characterized TnG repeat regions. Only the motifs significantly enriched (p<0.05) in FoxP3 ChIP-peaks over OCRs were analyzed. For FoxP3, 86.31% of H–T (red) and 84.41 % of T–H (blue) motif pairs overlap with TnG repeats. For FoxP1, 86.67% of H–T (green) and, 86.99% of T–H (brown) motif pairs overlap with TnG repeats.

The results reveal a strong enrichment of H-H pairs of G1-G1 motifs in FoxP3-bound regions across all three in vivo datasets (ChIP-seq, CNR-seq, and ChIP-exo; Figures 3B, S2A). The H-H motif pairs were found in 2,375 sites (2,270 unique peaks) within the total 26K FoxP3-ChIP peaks (184.96 per Mb), compared to 10,140 occurrences in 502,437 ATAC-peaks (80.34 per Mb). Among these 2,375 FoxP3-occupied H-H sites, the canonical IR-FKHM (inverted repeat of TGTTTNN, Figure 3B) appeared in 568 sites (24%), while the majority (76%) contained non-canonical motifs (i.e. non-IR-FKHM). FoxP3 preferentially bound H-H pairs of relaxed FKHM-like sequences (G1-G1) over G1-G2 or G2-G1, with G2-G2 pairings showing no significant enrichment (Figure 3B, 3C). This preference mirrored intrinsic affinity patterns observed in vitro using purified FoxP3 and DNA by EMSA (Figure 3D). Altogether, while IR-FKHM represents the most favored motif for FoxP3 in vivo and in vitro, non-canonical motifs collectively make substantial contributions to FoxP3’s H-H dimerization in cells.

In contrast to FoxP3, FoxP1 showed little sign of bias for H-H pairs both in vivo and in vitro. In CD8 T cells (chosen because they lack FoxP3 that can influence its genomic targeting), FoxP1 ChIP-seq data^20^ did not show any H-H motif enrichment (Figure 3B, S2B), consistent with our in vitro finding that FoxP1 does not display intrinsic preference for H-H motifs (Figure 1, 2).

In addition to the strong H-H enrichment, the second most prominent feature in the FoxP3 ChIP-seq heatmaps was the presence of H-T and T-H pairs of G1-G1 motifs (Figure 3B). We noted that TnG repeats––long tandem arrays of TnG or AnC sequences that are occupied by FoxP3 in vivo and in vitro^4,5,9^––were included in the H-T/T-H motif pair analysis (Figure 3E).

Specifically, 86.31% and 84.41% of significant H-T and T-H G1-G1 pairs, respectively, overlapped with TnG repeats within FoxP3 ChIP peaks (Figure 3F). By contrast, H-T/T-H arrangements of G2-G2 pairs were less enriched, likely because tandem repeats of GCAT (G2) and GCGT (G2- like), while capable of supporting FoxP3 multimerization (Figure S2C) and DNA bridging (Figures S2D-S2E), were less efficient than G1-G1 tandem repeats.

Note that these H-T/T-H pairs of G1-G1 motifs were also enriched in FoxP3 ChIP-exo and CNR- seq (Figure S2A), but were not enriched in oligo PD-seq (Figure 2B), presumably because the random-random oligo library only allowed 7+7 bp variation with fixed flanking and spacer sequences, which precluded TnG repeats. H-T/T-H G1–G1 pairs were also enriched in FoxP1 ChIP peaks (Figure 3B), where they too map primarily to TnG repeats (Figure 3F), consistent with the common ability of FoxP TFs to multimerize on these microsatellites^4^.

Overall, the paired motif analyses using genomic occupancy data indicate that FoxP3 employs both G1 and G2 motifs to form H-H dimers in vivo. In addition, FoxP3 forms H-T/T-H multimers on G1 motifs (TnG repeats). Although tandem repeats of G2-like sequences can also facilitate multimerization, they do so with markedly lower efficiency and enrichment. By contrast, T-T motifs were not enriched, suggesting that this configuration is not favored for FoxP3 dimerization or multimerization.

### 4. FoxP3-bound H-H motifs are enriched for a 4-nt gap and are preferentially associated with FoxP3-regulated genes

The observed enrichment of H-H motifs contrasts with a previous report^3^ that found no such enrichment. Given this contradiction, we further investigated the significance of the observed enrichment of H-H motifs. We first compared two analysis methodologies that led to the divergent outcomes. That earlier report^3^ was restricted to 5,047 ChIP peaks that are highly reproducible in both peak width and locations (defined by 50% reciprocal overlap across four ChIP-seq datasets^14,15^). In contrast, our analysis in Figure 3 used peaks directly called from the same four datasets with a threshold of *p*<0.001, yielding 26K peaks (see Methods for details). The H-H motif-containing loci showed concordant ChIP signal, regardless of the datasets (Figure S3A).

We first asked whether the broader peak definition might have artificially contributed to H-H enrichment, we compared gap-size distributions of significant motif pairs in FoxP3-bound loci (26K ChIP peaks) versus Treg OCRs (502K ATAC peaks). If enrichment were an artifact of peak broadening, no particular gap size should be favored. Indeed, OCRs showed no preference in gap size (Figure 4A), whereas FoxP3-bound regions were strongly enriched for a 4-nt gap across all significant motif pairs (Figure 4A)—the optimal spacing for H-H dimerization^3^. Similar enrichment was observed in FoxP3-bound loci defined by 19K CNR peaks (Figure S3B). These results indicate that H-H motif enrichment reflects genuine FoxP3 binding properties, not artifacts of peak definition.

**Figure 4.**
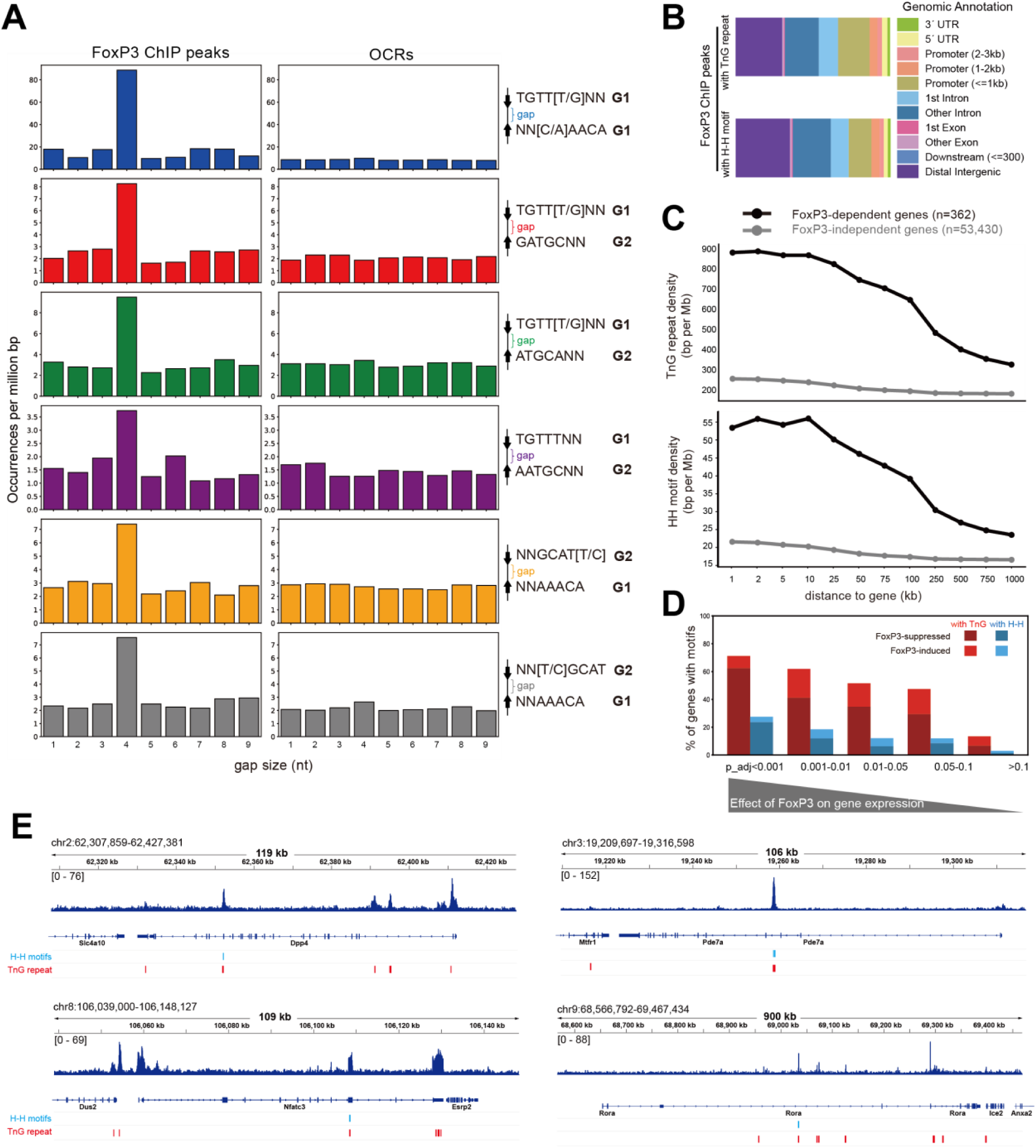
FoxP3-bound H-H motifs are enriched for a 4-nt gap and are preferentially associated with FoxP3-dependent genes. A. Gap size distributions for the indicated motif pairs across **FoxP3 ChIP-seq peaks** (left) and all **OCRs (**right). Motif frequency was calculated as in Figure 3A. Only the H-H motifs significantly enriched in FoxP3-bound sites in Figure 3A were analyzed. B. Genomic distribution of FoxP3 ChIP peaks with TnG repeats (top) and H-H motifs (bottom) in the *M. musculus* genome relative to Transcription Start Sites (TSSs). C. Quantification of TnG repeats (top) and H–H motifs (bottom) near FoxP3-dependent genes. Genes were stratified into FoxP3-dependent (adjusted p-value < 0.05, black, n=362) and FoxP3-independent (adjusted p-value ≥ 0.05, gray, n=53,430) groups based on RNA-seq differential expression following acute FoxP3-degradation in activated Treg (PMID:40478934). For each gene, regions extending from 1 kb to 1 Mb were analyzed, and the density of overlapping TnG repeats or H–H motifs within these regions was calculated. The plots indicate averaged values among genes in each group. D. Fraction of FoxP3-dependent genes harboring TnG (red) or H-H motifs (blue) within 2 kb expansion from each gene. Genes were binned by adjusted p-value from the differential expression analysis (see above) and further classified as FoxP3-suppressed vs. -induced groups. A higher fraction of FoxP3-regulated genes contained nearby TnG or H–H motifs within 2 kb, with the majority of these genes suppressed rather than induced by FoxP3. E. Genome browser views of representative FoxP3-dependent genes containing H–H motifs and TnG repeats. Shown are FoxP3 ChIP-seq tracks with motif locations for Dpp4 (top left), Pde7a (top right), Nfatc3 (bottom left), and Rora (bottom right).

Why, then, was H-H motif enrichment missed in the earlier analysis^3^? Although most of the earlier 5K peaks (4,738) overlapped with our current 26K peak set, the smaller dataset may have lacked sufficient statistical power. Additionally, the narrow focus on the strict IR-FKHM rather than broader H-H motifs could have further contributed. Consistent with both of these ideas, re-analysis of the original 5K peaks with our expanded H-H definitions revealed that, while a modest 4-nt gap preference was observed for some non-canonical motifs (e.g., TGTTKNN_GATGCNN and NNGCATY_NNAAACA) (Figure S3C), the baseline distribution was uneven, diminishing statistical robustness. Together, these findings suggest that restrictive motif definitions combined with limited peak sets led to the failure of the earlier study^3^ to detect H-H enrichment.

To evaluate the functional significance of H-H motifs and compare with that of TnG repeats, we analyzed their genomic distribution. Within the 26K FoxP3 ChIP peaks, both H-H motifs and TnG repeats were broadly distributed across various genomic features, from promoter-proximal regions to gene-distal sites (Figure 4B). Histone mark analyses further supported this widespread distribution (Figure S3D). To link motif binding to gene regulation, we focused on genes affected by acute FoxP3 degradation^21^, as FoxP3 knockout models are known to alter hundreds of genes indirectly^14,22^. Strikingly, genes directly regulated by FoxP3 contained significantly higher densities of both H-H motifs and TnG repeats than unaffected genes (Figure 4C).

We further examined the fraction of FoxP3-regulated genes that harbor H-H motifs or TnG repeats within 2 kb. Based on differential gene expression analysis, genes most significantly affected by FoxP3 depletion (lower p-values) were far more likely to contain these motifs, whereas this enrichment declined with the significance of FoxP3’ effect (Figure 4D).

Importantly, most motif-associated, FoxP3-regulated genes were upregulated upon FoxP3 loss, consistent with its predominant role as a suppressor^21^. Many of these FoxP3-regulated genes encode factors involved in T cell biology (e.g., *Rora, Nfatc3, Pde7a*, and *Dpp4*) (Figure 4E). Thus, these results support a model in which FoxP3 binds H-H motifs and TnG repeats to finetune Treg transcriptional program.

### 5. H-H dimerization seeds FoxP3 multimerization on adjacent TnG repeats

Our analyses above suggest that although TnG repeats are more frequent than H-H motifs, the two often occur in close proximity and share similar genomic features, raising the question of how FoxP3 dimerization on H-H motifs relates to its multimerization on TnG repeats. A systematic analysis of FoxP3 ChIP peaks revealed that 68.9% of H-H–containing peaks (1,563 of 2,270) also harbored TnG repeats (Figure 5A). By contrast, only 26.9% of H-H–containing open chromatin regions (2,697 of 10,027 ATAC peaks) overlapped with TnG repeats (Figure 5A).

**Figure 5.**
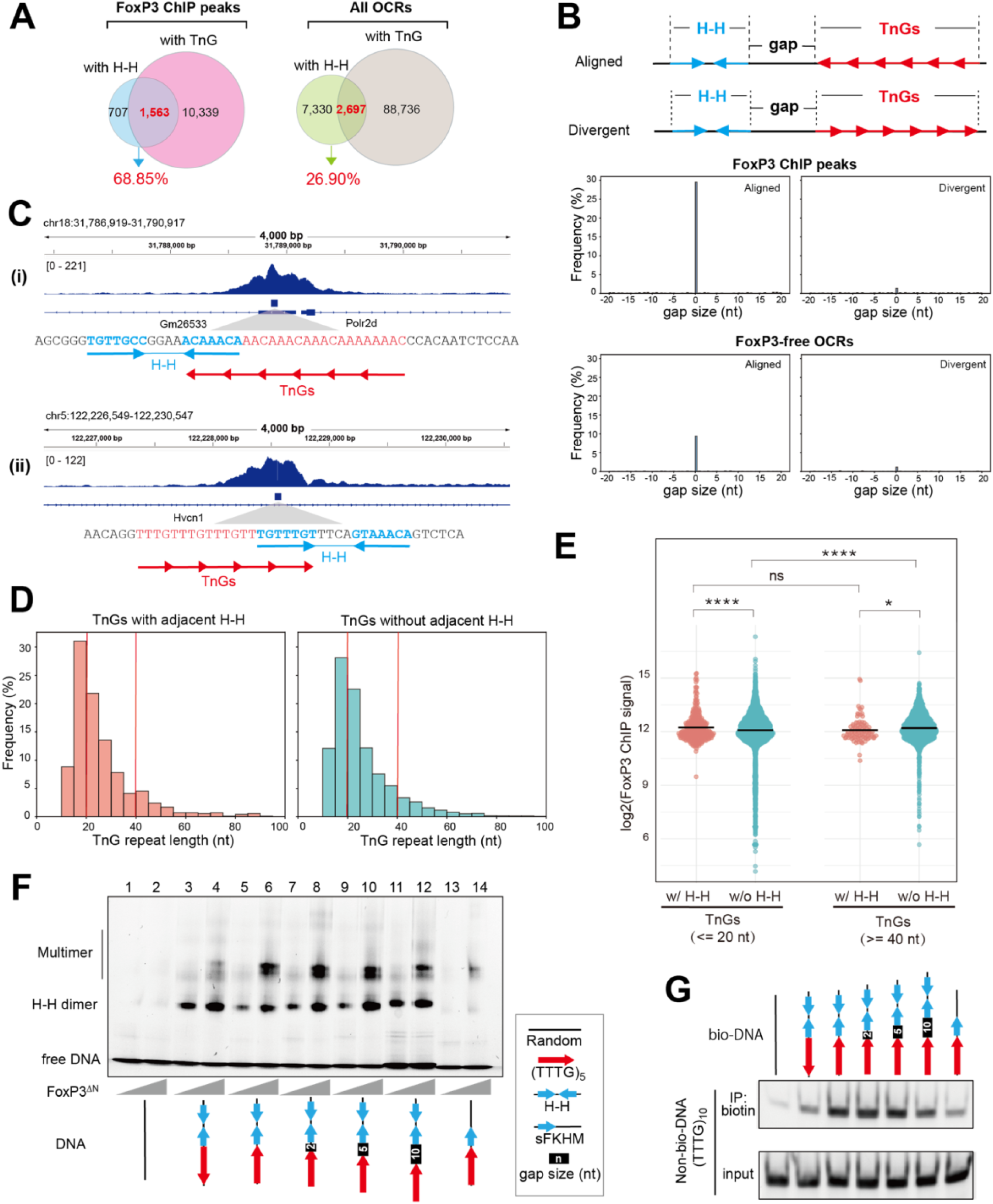
H-H dimerization seeds FoxP3 multimerization on adjacent TnG repeats. A. Overlap between FoxP3 ChIP peaks (or all OCRs) containing H-H motif pairs vs. TnG repeats. 68.9% of H–H-containing FoxP3 peaks harbor TnG repeats, whereas only 26.9% of H-H- containing OCRs harbor TnG repeats. B. Top: Two possible configurations of H-H motifs and TnG repeats. A TnG repeat and the proximal half-site of the H-H motif can be either "aligned" or "divergent." Bottom: Distribution of distances between the borders of H-H motifs and the nearest TnG repeat in the aligned (left) and divergent (right) configurations. We measured the border-to-border distance as positive when the TnG occurred downstream, and negative when it occurred upstream. C. Genome browser views of Foxp3 ChIP peaks harboring TnG repeats with adjacent H-H with 0 nt gap in the aligned configuration. Tracks (top to bottom): Foxp3 ChIP-seq, TnG repeats with adjacent H-H within Foxp3 Chip-seq peaks, and Refseq gene annotation. D. Histogram analysis of TnG repeat lengths, comparing TnG repeats with 0 nt gap H–H motifs in the aligned configuration (left, n=702) vs. those that are more than 1 kb away from H–H motif (right, n=16,144). Red vertical lines indicate length thresholds used in (E). E. Dot plots display the log2-transformed FoxP3 ChIP-seq signal (measured as the area under the curve [AUC] within ±100 nt from the center of TnG repeats) for two groups: TnG repeats <=20 nt or >=40 nt. In each group, TnG repeats with (pink) and without (teal) adjacent to H– H motifs (as defined in D) were compared. p < 0.0001 (****), p < 0.05 (*), p> 0.01 (ns) using two-way ANOVA with Tukey’s multiple comparisons test. F. Native gel shift assay of MBP-tagged FoxP3^ΔN^ (0.4 or 0.8 μM) with DNAs (0.2 μM) containing TTTG repeats and IR-FKHM (H-H) with a 0-10 nt gap (see Table S1 for sequences). G. DNA bridging assay. Biotinylated DNA (bio-DNA, 60 bp) of indicated sequence and non- biotinylated DNA of (TTTG)_10_ (40 bp) were mixed at a 1:1 ratio (0.1 μM each), incubated with FoxP3^ΔN^ (0.4 μM) and processed for Streptavidin pull-down as in Figure S2D. Non- biotinylated DNA in the eluate was analyzed on the gel (SybrGold).

To better define the spatial relationship between H-H motifs and TnG repeats, we measured distances between these motifs (border-to-border distance), designating positive values when a TnG repeat occurred downstream and negative values when upstream (Figure 5B, top).

Overlaps were counted as a 0 nt gap. This analysis revealed a pronounced peak at 0 nt in FoxP3- bound loci (Figure 5B, bottom; examples in 5C), with 29.6% of H-H sites (702 of 2,375) lying immediately adjacent to or overlap with a TnG repeat—compared with only 9.4% in FoxP3-free OCRs. Strikingly, these abutting motifs displayed a strong orientational bias, such that the proximal half-site of the H-H motif could seamlessly extend into the neighboring TnG repeat (“aligned”; Figure 5B, 5C).

Prompted by this close positioning, we asked whether H-H proximity enhances FoxP3 occupancy on TnG repeats. We compared ChIP signals at repeats with adjacent H-H motifs (0 nt gap) to those located > 1 kb from any H-H site. We stratified TnG repeats into short (≤ 20 bp) and long (≥ 40 bp) groups (Figure 5D), because longer repeats inherently bind FoxP3 more efficiently^4^. Although H-H adjacency boosted FoxP3 occupancy in both groups, the effect was markedly greater for short repeats (Figure 5E). As a result, TnG repeats adjacent to H-H motifs exhibited little dependence on the repeat length, whereas TnG repeats without H-H showed FoxP3 occupancy that increased with length. These findings indicate that H–H dimerization seeds or stabilizes FoxP3 multimerization on otherwise suboptimal, short TnG repeats.

In support of this notion, 5 repeat of TTTG [(TTTG)_5_]––which alone is too short to drive robust FoxP3 multimerization^4^, even with a single adjacent FKHM (lanes 13-14, Figure 5F)––greatly facilitated multimer assembly when an IR-FKHM was placed immediately adjacent (lanes 5-6, Figure 5F). Moreover, this positive effect was seen only when IR-FKHM was in the “aligned” orientation as (TTTG)_5_ and with a 0-5 nt-gap, but not in the “divergent” configuration (lanes 3-4) or with a 10 nt-gap (lanes 5-12). Likewise, DNA-bridging on (TTTG)_5_ was substantially enhanced by a neighboring H-H motif only in the aligned orientation with a 0-5 nt-gap (Figure 5G).

Together, these findings demonstrate that H-H dimerization expands FoxP3’ TnG sequence specificity, enabling efficient multimerization and DNA bridging even on otherwise suboptimal, short TnG repeats.

### 6. RBR is the key determinant for H-H dimerization of FoxP3

Our finding that only FoxP3, and not FoxP1, can form H-H dimer, while both can form H-T multimers, raises the question regarding the molecular basis of this specificity. It is particularly intriguing given that FoxP3 and FoxP1 share conserved domain architectures and similar FKH DNA-binding domains (Figure 1A, 6A). We first investigated whether a small number of divergent residues within the FKH DBD (e.g. R358, N376, H377, P378, S400, E401, and R414, Figure S4A, S4B) underlie FoxP3’s unique H-H dimerization. However, mutating these residues minimally affected FoxP3’s preference for IR-FKHM over sFKHM (Figure S4C), suggesting that they are not the primary determinants of FoxP3’s capacity for H-H dimer formation.

Previous studies showed that the hydrophobic loop (namely Runx1-binding region, RBR) immediately preceding FKH DBD is important for both H-H dimerization and H-T multimerization of FoxP3^3–5^. Notably, the sequence of FoxP3’s RBR differs considerably from that of FoxP1, FoxP2, and FoxP4 (Figure 6A). We therefore investigated whether RBR alone dictates the divergent ability to form H-H dimer by FoxP3 and FoxP1. Note that removal of RBR leads to misfolding of FKH of all four FoxP TFs, from the winged-helix conformation to domain-swap dimer^3^, precluding RBR deletion test. We instead swapped the RBR between FoxP3 and FoxP1 and examined the impact on their DNA binding behaviors (Figure 6B). We used RBR-FKH constructs (FoxP^RBR-FKH^)—the minimal constructs required for proper folding and maintenance of full-length protein’s DNA specificity^3^. Unlike the FoxP^ΔN^ constructs, which forms a constitutive dimer through a coiled-coil (CC) domain^13^, FoxP^RBR-FKH^ is monomeric^3^, allowing us to distinguish monomeric from dimeric binding based on distinct EMSA band patterns. Since monomeric binding is generally weak^3^, NFAT was used to stabilize FoxP3–DNA interactions, enabling reliable capture of both monomeric and dimeric complexes.

**Figure 6.**
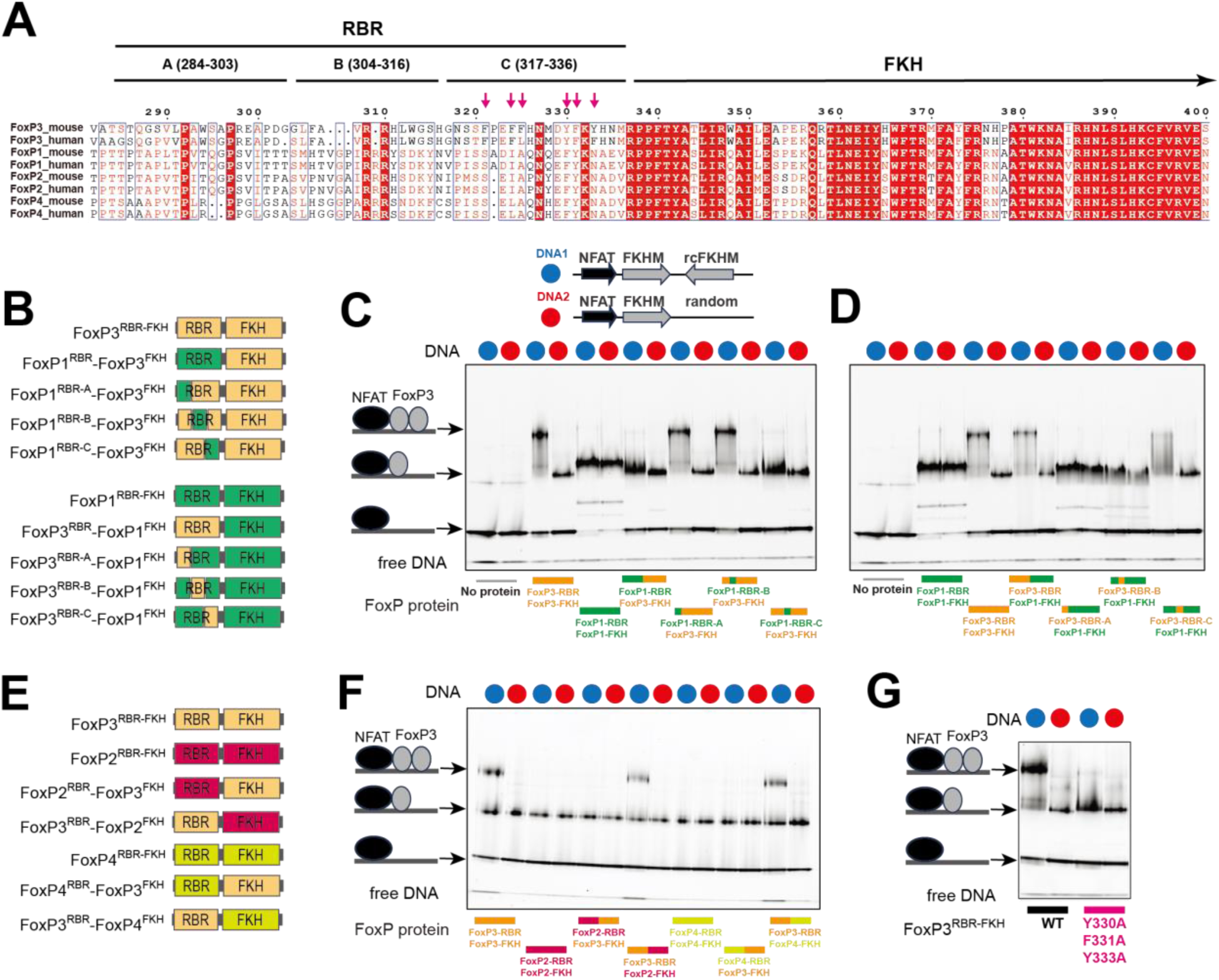
RBR is the key determinant for H-H dimerization of FoxP3. A. Sequence alignment of mouse and human FoxP TFs, with moderately conserved residues indicated in red font and highly conserved residues highlighted in red. The FKH domain is well conserved across the family, whereas the RBR loop region of FoxP3 is unique compared to FoxP1, FoxP2, and FoxP4. For chimeric construct design, the RBR loop is divided into three segments (A, B, and C). Arrows indicate six aromatic residues within the RBR-C loop of mouse FoxP3. B. Chimeric constructs involving full or partial RBR loop swaps between FoxP3 and FoxP1. FoxP1 sequences are labeled in green, FoxP3 sequences are labeled in yellow. RBR was divided into three segments (as in panel **A**) for partial swaps. C-D. Native gel shift assay of various chimeric RBR-FKH constructs (0.8 μM) with DNA (0.2 μM) harboring IR-FKHM (blue) or sFKHM (red) in the presence of NFAT (0.4 μM). FoxP3^RBR-FKH^ binds to DNA with IR-FKHM (DNA1, blue) as a dimer, whereas it binds to DNA with single FKHM (DNA2, red) as a monomer. In contrast, FoxP1^RBR-FKH^ binds both DNA 1 and DNA2 as a monomer. This difference between FoxP3 and FoxP1 was entirely dependent on RBR-C, as shown by the RBR and RBR-C swaps. E. Chimeric constructs involving full RBR loop-swaps between FoxP3 and FoxP2/FoxP4. FoxP2, FoxP3 and FoxP4 sequences are labeled in pink, yellow, and light green, respectively. F. Native gel shift assay of various chimeric RBR-FKH constructs (0.8 μM) incubated with DNA (0.2 μM) harboring IR-FKHM (blue) or sFKHM (red) in the presence of NFAT (0.4 μM). Only constructs with the FoxP3 RBR can bind IR-FKHM DNA as a dimer. All other constructs bind both sFKHM and IR-FKHM as a monomer, suggesting the importance of FoxP3 RBR in H-H dimerization. G. Effect of the aromatic mutations (Y330A/F331A/Y333A) on FoxP3 dimerization on IR-FKHM DNA. FoxP3^RBR-FKH^ (0.8 μM) with or without mutations were incubated with IR-FKHM or sFKHM DNA (0.2 μM) in the presence of NFAT (0.4 μM). Only the wild-type forms a dimer on IR-FKHM, suggesting the importance of the aromatic residues in H-H dimerization.

Consistent with a previous report^3^, FoxP3^RBR-FKH^ bound IR-FKHM as a dimer, but single isolated FKHM (sFKHM) as a monomer (Figure 6C, 6D). In contrast, FoxP1^RBR-FKH^ bound both IR-FKHM and sFKHM as a monomer (Figure 6C, 6D), even when protein was in excess over DNA. This indicates that FoxP1 is inherently incompatible with double occupancy on IR-FKHM. Remarkably, swapping the RBRs interchanged their binding modes. That is, a chimera with FoxP1^RBR^ and FoxP3^FKH^ bound DNA in the same manner as FoxP1, not FoxP3 (Figure 6C).

Conversely, a chimera comprising the FoxP3^RBR^ and FoxP1^FKH^ bound DNA in the same manner as FoxP3, not FoxP1 (Figure 6D). Furthermore, a similar shift in binding mode was observed upon exchanging RBR loops between FoxP3 and either FoxP2 or FoxP4 (Figure 6E, 6F). When swapping FoxP3 RBR loop with that of FoxP3 from different species, we preserved dimeric binding modes (Figure S4D, S4E), although with a modest reduction in DNA binding affinity for some RBR species. Collectively, these findings indicate that RBR is a modular element and the sole determinant for H-H dimerization in FoxP3.

To further delineate the specific region within the RBR loop responsible for FoxP3’s distinct H-H dimerization property, we divided the loop into three segments (A, B, and C, Figure 6A), each comprising 20 amino acids. Through targeted sequence-swapping, we identified fragment C (amino acids 317-336; RBR-C) as critical for FoxP3’s H-H dimerization. Substituting FoxP3’s RBR- C with the corresponding region from FoxP1 abolished FoxP3’s ability to form dimers on the IR-FKHM element, whereas introducing FoxP3’s RBR-C into FoxP1 conferred dimerization capacity (Figure 6C, 6D).

A comparison of RBR-C sequences revealed that FoxP3 contains a higher density of aromatic residues than that of FoxP1/2/4: six out of 20 residues in FoxP3’s RBR-C are aromatic, compared with only two in FoxP1/2/4 (Figure 6A). Mutating these aromatic residues (Y330A, F331A, Y333A) significantly reduced FoxP3’s ability to bind IR-FKHM as a dimer (Figure 6G). These aromatic residues also contribute to FoxP3’s multimerization on TnG repeat DNA and FoxP3’s transcriptional activity in CD4 T cells^5^, underscoring the importance and the dual role of these aromatic RBRs in mediating both H-H dimerization and multimerization.

In addition to differences in aromatic content, secondary structure predictions indicated an α- helix in the RBR-C region of FoxP1, whereas FoxP3 was predicted to contain only a short, four– amino acid helical stretch (Figure S4F). NMR chemical shifts confirmed a strong α-helical propensity for FoxP1 residues 477-488 within RBR-C (Figure S4G). In case of FoxP3, NMR signals were weaker and displayed line broadening––indicative of conformational exchange occurring on the NMR timescale––which precluded direct comparison between FoxP1 and FoxP3 RBR conformations. Nonetheless, the previously reported structures of FoxP3 H-H dimer^3^ and H-T multimers^4,5^ revealed both α-helical and non-helical, disordered RBC-C conformations, consistent with its intrinsically lower helical propensity.

Taken together, these results pinpoint RBR-C as the primary driver of H-H dimerization in FoxP3, possibly due to differences in hydrophobicity and helical propensity.

## DISCUSSION

Previous studies have shown that FoxP3 binds DNA in two distinct modes: a multimeric conformation when binding long (>30 bp) TnG repeats (n = 2-5)^4,5^ and a head-to-head (H-H) dimeric conformation on shorter motifs such as IR-FKHM^3^. FoxP3 multimerization occurs at numerous sites, facilitating chromatin looping through DNA bridging and global transcriptional regulation^4,5^ (Figure 7A). In contrast, in vivo evidence for—and the functions of—FoxP3’s H-H dimerization has remained elusive. Using a degenerate oligonucleotide pull-down approach, we identified a broader repertoire of sequences that support H-H binding (Figure 7B). Both FKHM- like (group 1) and GCAT-containing motifs (group 2) can support FoxP3 binding in H-H orientation with a preferred 4-nt gap. These motifs were enriched not only in vitro but also at FoxP3-occupied sites in vivo, as revealed by ChIP-seq, ChIP-exo, and CNR-seq. Their enrichment was overlooked in our earlier analysis^3^ due to a narrow focus on IR-FKHM and the use of a limited number of ChIP peaks, which reduced statistical power. Importantly, both H-H motifs and TnG repeats were more frequently associated with FoxP3-regulated genes than FoxP3- independent genes in Tregs, underscoring their functional relevance.

**Figure 7.**
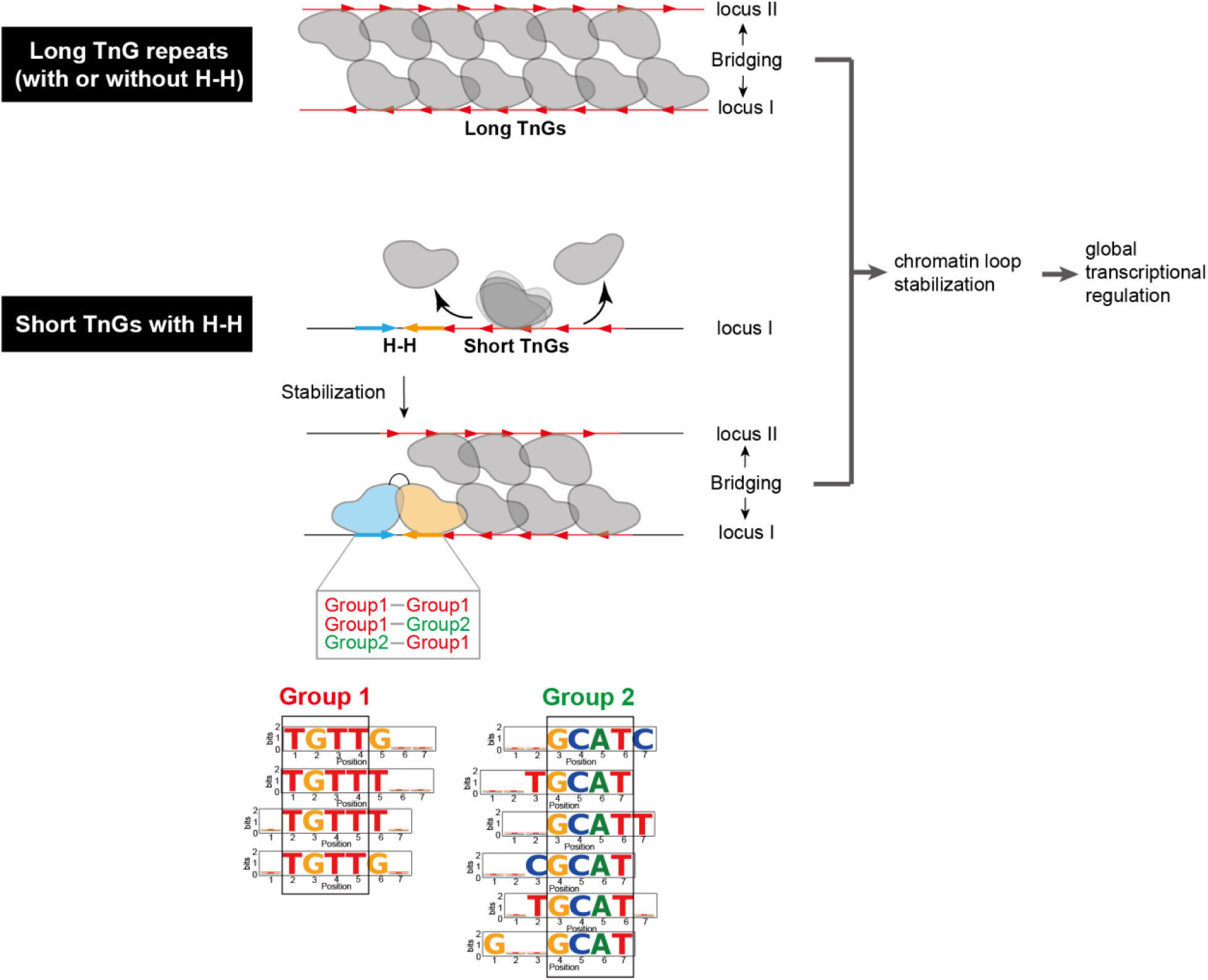
Role of FoxP3 H-H dimerization in gene regulation. FoxP3 bridges genomic loci and regulate global transcriptome of Treg by multimerizing on TnG repeat microsatellites. In addition, FoxP3 can form head-to-head (H-H) dimers in Tregs. Although less frequent, dimerization likely plays an important role by stabilizing FoxP3 multimers. On long TnG repeats, FoxP3 forms stable multimers that bridge distinct DNA segments independently of H-H motifs. By contrast, on shorter or suboptimal TnG repeats, FoxP3 multimerization is unstable, and may require adjacent H-H motifs for stabilization. Both Group 1 (FKHM-like sequences) and Group 2 sequences can support H-H dimer formation. In summary, H-H dimerization alone does not support DNA bridging but functions to extend FoxP3’s architectural activity to weaker TnG repeats.

What, then, are the molecular functions of FoxP3’s two binding modes? Our analyses showed that TnG repeats are more common (∼50% of ChIP peaks) compared with H-H motifs (∼10%). Despite their low frequency, H-H motifs often reside adjacent to TnG repeats, where they can “seed” or stabilize FoxP3 multimerization on neigthboring sequences, particularly on shorter or suboptimal repeats that would otherwise fail to form stable multimers (Figure 7B). Once established, FoxP3 multimerization promotes DNA bridging and stabilization of chromatin loops^4,5^, enabling global transcriptional regulation––either directly or indirectly––through modulation of chromatin architecture. This model is in keeping with the finding that FoxP3 typically exerts only modest effects (<2–3 fold) across hundreds of genes, with its regulatory impact highly context-dependent^21^. We therefore propose that H-H motifs serve a complementary role to TnG repeats, extending FoxP3’s architectural influence to a wider genomic repertoire and thereby broadening its ability to maintain the Treg transcriptional landscape.

## ACKNOWLEDGEMENT

We thank all members of the Hur lab for their helpful discussion and feedback. This study was supported by NIH grants (R01AI180137 to S.H.) and Howard Hughes Medical Institute (S.H.).

## STAR METHODS

### KEY RESOURCES TABLE

| REAGENT or RESOURCE | SOURCE | IDENTIFIER |
| --- | --- | --- |
| **Chemicals, Peptides, and Recombinant Proteins** |  |  |
| Ni-NTA agarose | QIAGEN | Cat# 30250 |
| Amylose Resin | New England Biolabs | Cat# E8021L |
| HRV 3C protease | Homemade | N/A |
| HiTrap Heparin | GE Healthcare | Cat# 17040601 |
| Superdex 200 Increase 10/300 | GE Healthcare | Cat# 28990944 |
| Superdex 75 Increase 10/300 | GE Healthcare | Cat# 29148721 |
| HiTrap SP | GE Healthcare | Cat# 17505301 |
| High Capacity Streptavidin Agarose | Thermo Fisher | Cat# 20359 |
| SYBR Gold stain | Invitrogen | Cat# S11494 |
| Phusion polymerase | New England Biolabs | Cat# M0530L |
| DNA Polymerase I, large fragment | New England Biolabs | Cat# M0210S |
| XbaI | New England Biolabs | Cat# R0145S |
| BamHI-HF | New England Biolabs | Cat# R3136 |
| XmaI | New England Biolabs | Cat# R0180S |
| HindIII-HF | New England Biolabs | Cat# R3104S |
| Deoxyribonuclease | Sigma | Cat# DN25 |
| Bis-Tris native PAGE, 3-12% | Invitrogen | Cat# BN1003BOX |
| Novex TBE Gels, 10% | Invitrogen | Cat# EC62755BOX |
| Mini-PROTEAN TGX Precast Protein Gels, 4-15% | BioRad | Cat# 4561086 |
| Uranyl formate | Electron Microscopy Sciences | Cat# 22451 |
| Glutaraldehyde solution | Sigma | Cat# G5882 |
| TriDye™ Ultra Low Range DNA Ladder | New England Biolabs | Cat# N0558S |
| **Critical Commercial Assays** |  |  |
| QIAquick Nucleotide Removal kit | QIAGEN | Cat# 28304 |
| **Deposited Data** |  |  |
| NMR | This study | BMRB ID: 5301 |
| **Experimental Models: Organisms/Strains** |  |  |
| *E. coli* BL21 (DE3) | Stratagene | Cat# 230130 |
| **Oligonucleotides** |  |  |
| DNAs | See Table S1 | N/A |
| Biotin-DNAs | See Table S1 | N/A |
| **Recombinant DNA** |  |  |
| pET50b-mFoxP3<sup>ΔN</sup> | This study | N/A |
| PMAL-mFoxP3<sup>ΔN</sup> | This study | N/A |
| PMAL-mFoxP1<sup>ΔN</sup> | This study | N/A |
| pET50b-mFoxP3<sup>RBR+FKH</sup> | This study | N/A |
| pET50b-mFoxP1<sup>RBR+FKH</sup> | This study | N/A |
| pET50b-mFoxP2<sup>RBR+FKH</sup> | This study | N/A |
| pET50b-mFoxP4<sup>RBR+FKH</sup> | This study | N/A |
| pET50b-mFoxP1<sup>RBR</sup>-FoxP3<sup>FKH</sup> | This study | N/A |
| pET50b-mFoxP2<sup>RBR</sup>-FoxP3<sup>FKH</sup> | This study | N/A |
| pET50b-mFoxP4<sup>RBR</sup>-FoxP3<sup>FKH</sup> | This study | N/A |
| pET50b-mFoxP3<sup>RBR</sup>-FoxP1<sup>FKH</sup> | This study | N/A |
| pET50b-mFoxP3<sup>RBR</sup>-FoxP2<sup>FKH</sup> | This study | N/A |
| pET50b-mFoxP3<sup>RBR</sup>-FoxP4<sup>FKH</sup> | This study | N/A |
| pET50b-mFoxP1<sup>RBR-A</sup>-FoxP3<sup>FKH</sup> | This study | N/A |
| pET50b-mFoxP1<sup>RBR-B</sup>-FoxP3<sup>FKH</sup> | This study | N/A |
| pET50b-mFoxP1<sup>RBR-C</sup>-FoxP3<sup>FKH</sup> | This study | N/A |
| pET50b-mFoxP3<sup>RBR-A</sup>-FoxP1<sup>FKH</sup> | This study | N/A |
| pET50b-mFoxP3<sup>RBR-B</sup>-FoxP1<sup>FKH</sup> | This study | N/A |
| pET50b-mFoxP3<sup>RBR-C</sup>-FoxP1<sup>FKH</sup> | This study | N/A |
| pET50b-mFoxP3<sup>RBR+FKH</sup>-Y330A/F331A/Y333A | This study | N/A |
| pET50b-mFoxP3<sup>RBR+FKH</sup>-N376R/H377N/P378A | This study | N/A |
| pET50b-mFoxP3<sup>RBR+FKH</sup>-S400N/E401V | This study | N/A |
| pET50b-mFoxP3<sup>RBR+FKH</sup>-R414Q | This study | N/A |
| pET50b-FoxP3<sup>RBR+FKH</sup>-Platypus | This study | N/A |
| pET50b-FoxP3<sup>RBR+FKH</sup>-Opossum | This study | N/A |
| pET50b-FoxP3<sup>RBR+FKH</sup>-Frog | This study | N/A |
| pET50b-FoxP3<sup>RBR+FKH</sup>-Zebrafish | This study | N/A |
| pET50b-FoxP3<sup>RBR-C</sup>-Platypus-mFoxP3<sup>FKH</sup> | This study | N/A |
| pET50b-FoxP3<sup>RBR-C</sup>-Opossum-mFoxP3<sup>FKH</sup> | This study | N/A |
| pET50b-FoxP3<sup>RBR-C</sup>-Frog-mFoxP3<sup>FKH</sup> | This study | N/A |
| pET50b-FoxP3<sup>RBR-C</sup>-Zebrafish-mFoxP3<sup>FKH</sup> | This study | N/A |
| **Software and Algorithms** |  |  |
| Eman2<sup>1</sup> | G Tang et al., 2007 | https://blake.bcm.edu/emanwiki/EMAN2 |
| BBMerge (v38.84)<sup>2</sup> | Bushnell B et al., 2017 | https://jgi.doe.gov/data-and-tools/bbtools/bb-tools-user-guide/bbmerge-guide/ |
| Biopython (v1.85)<sup>3</sup> | Cock et al., 2009 | https://biopython.org/ |
| Pandas (v2.2.3)<sup>4</sup> | McKinney W, 2010 | https://pandas.pydata.org |
| Matplotlib (v3.9.4)<sup>5</sup> | Hunter JD, 2007 | https://matplotlib.org/ |
| Numpy (v2.0.2)<sup>6</sup> | Harris CR et al., 2020 | https://numpy.org/ |
| Bowtie2 (v2.3.4.3)<sup>7</sup> | Langmead and Salzberg, 2012 | http://bowtie-bio.sourceforge.net/bowtie2/index.shtml |
| SAMtools (v1.9)<sup>8</sup> | Danecek P et al., 2021 | http://www.htslib.org/ |
| MACS3 (v3.0.3)<sup>9</sup> | Zhang et al., 2008 | https://github.com/macs3-project/MACS |
| Bedtools (v2.31.0)<sup>10</sup> | Aaron R et al., 2010 | https://bedtools.readthedocs.io/en/latest/ |
| SRATools (v3.1.1)<sup>11</sup> | Leinonen R et al., 2011 | https://github.com/ncbi/sra-tools |
| Trimmomatic (v0.36)<sup>12</sup> | Bolger AM et al., 2014 | http://www.usadellab.org/cms/?page=trimmomatic |
| FIMO (v5.5.8)<sup>13</sup> | Grant CE et al., 2011 | https://meme-suite.org/meme/doc/fimo.html |
| MEME-ChIP (v5.3.3)<sup>14</sup> | Bailey TL et al., 2015 | https://meme-suite.org/meme/tools/meme-chip |
| Pyranges (v0.1.4)<sup>15</sup> | Stovner EB et al., 2020 | https://github.com/biocore-ntnu/pyranges |
| ChIPseeker (v1.42.1)<sup>16</sup> | Kitagawa Y et al., 2017 | https://bioconductor.org/packages/ChIPseeker |
| DeepTools (v3.5.6)<sup>17</sup> | Ramirez F et al., 2016 | https://deeptools.readthedocs.io/en/develop/ |
| Analysis codes | This study | <u>[DOI: 10.5281/zenodo.1710850](https://doi.org/10.5281/zenodo.1710850)</u><br><u>1</u> |

## EXPERIMENTAL MODEL AND STUDY PARTICIPANT DETAILS METHOD DETAILS

### Material Preparation

#### Plasmids

For bacterial expression, the genes encoding mouse FoxP3ΔN (residues 188-423) and mouse FoxP1 ΔN (residues 342-579) were inserted into pET50b between Xmal and HindIII sites, and into a pMAL-c2 vector between BamHI and Xbal sites respectively. The pMAL-c2 vector was modified to contain a HRV 3C protease target sequence between EcoRI and BamHI sites. Other constructs including mouse FoxP3RBR-forkhead (residues 284-423), mouse FoxP1RBR-forkhead (residues 427-423), mouse FoxP2RBR-forkhead (residues 447-588), mouse FoxP4RBR-forkhead (residues 418-557), FoxP3RBR-forkhead from different species, and chimeric constructs were generated by inserting genes into pET50b between Xmal and HindIII sites. All FoxP3 mutations including Y330A/F331A/Y333A, N376R/H377N/P378A, S400N/E401V and R414Q were generated by site-directed mutagenesis using Phusion High Fidelity (New England Biolabs) DNA polymerases.

#### DNA oligos

Single-stranded DNA (ssDNA) oligonucleotides were synthesized by IDT (Integrated DNA Technologies). To generate the degenerate DNA library for FoxP3 and FoxP1 PD-seq, 30 µL of ssDNA (200 µM) was annealed with 60 µL of a complementary primer (Klenow-1, 200 µM) in 10 µL 10× NEB Buffer 2 (final volume: 100 µL). Annealing was performed by heating to 95°C for 5 min followed by slow cooling to room temperature. The reaction was supplemented with 10 µL of 10 mM dNTP mix and 2 µL of Klenow fragment (DNA Polymerase I, large fragment; NEB), and incubated at 25°C for 30–60 minutes to synthesize the complementary strand. The resulting dsDNA was purified using the QIAquick Nucleotide Removal Kit (Qiagen). DNA purity and integrity assessed by TBE gel electrophoresis and Agilent TapeStation analysis. To prepare biotin-labeled dsDNA for DNA bridging assays, long forward ssDNA was mixed with a short biotinylated complementary ssDNA at equimolar concentrations. After annealing, the Klenow fragment was used to extend the complementary strand, generating full-length biotin-labeled dsDNA. Double-stranded DNA (dsDNA) oligos for EMSA assay, pulldown assay and DNA bridging assay were annealed from single-stranded, complementary oligos. Sequence of all the DNA oligos used are shown in Table S1.

#### Protein expression and purification

All recombinant proteins in this paper were expressed in BL21(DE3) at 18°C for 16-20 hrs following induction with 0.2 mM IPTG. Cells were lysed by high-pressure homogenization using an Emulsiflex C3 (Avestin). All proteins are from the Mus. musculus sequence, unless mentioned otherwise. His6-MBP-tagged FoxP3ΔN (residues 188-423) and FoxP1 ΔN (residues 342-579) used for PD-seq and DNA PD were purified using Ni-NTA affinity column and Superdex 200 Increase 10/300 (GE Healthcare) SEC column in 20 mM Tris-HCl pH 7.5, 500 mM NaCl, 2 mM DTT. FoxP3ΔN (residues 188-423) was expressed as a fusion protein with an N-terminal His6-NusA tag. After purification using Ni-NTA agarose, the protein was treated with HRV3C protease to cleave the His6-NusA-tag and were further purified by a series of chromatography purification using HiTrap Heparin (GE Healthcare), Hitrip SP (GE Healthcare) and Superdex 200 Increase 10/300 (GE Healthcare) columns. The final size-exclusion chromatography (SEC) was done in 20 mM Tris-HCl pH 7.5, 500 mM NaCl, 2 mM DTT. NFAT1 protein (residues 394-680) was also expressed as a fusion protein with an N-terminal His6-NusA tag. After purification using Ni-NTA agarose, the His6-NusA-tag was removed using the HRV3C protease and was further purified by SEC on Superdex 75 Increase 10/300 (GE Healthcare) column in 20 mM Tris-HCl pH 7.5, 500 mM NaCl, 5% Glycerol, 2 mM DTT. His6-NusA tagged proteins are all purified using Ni-NTA affinity column and Superdex 200 Increase 10/300 (GE Healthcare) SEC column in 20 mM Tris-HCl pH 7.5, 500 mM NaCl, 2 mM DTT. All RBR-forkhead proteins (including all chimeric proteins) were expressed as a fusion protein with an N-terminal His6-NusA tag. After purification using Ni-NTA agarose, the protein was treated with HRV3C protease to cleave the His6-NusA-tag and were further purified by Superdex 75 Increase 10/300 (GE Healthcare) column in 20 mM Tris-HCl pH 7.5, 500 mM NaCl, 2 mM DTT.

#### DNA pulldown assay using MBP tagged proteins

0.4 μM purified MBP-mFoxP3ΔN or MBP-mFoxP1ΔN protein was incubated with 0.1 μM DNA in the incubation buffer (20 mM Tris-HCl pH 7.5, 100 mM NaCl, 1.5 mM MgCl2) for 20 mins. The protein-DNA mixture was then incubated with 25μL Amylose Resin (New England Biolabs) for 30 mins with rotation at room temperature. The bound DNA was recovered using proteinase K (New England Biolabs), purified using QIAquick Nucleotide Removal kit (QIAGEN) and analyzed on 10% Novex TBE Gels (Invitrogen). DNA was visualized by SYBR Gold staining. For NFAT–FoxP3 cooperativity assay, 0.2 μM NFAT protein (residues 394–680) was included along with 0.4 μM MBP-mFoxP3ΔN and 0.1 μM DNA. All other steps were performed as described above.

#### DNA bridging assay

Biotin-DNA (bait, 0.1μM) was incubated with Streptavidin Agarose (25μL, Thermo Fisher) in buffer B (20mM Tris-HCl pH 7.5, 100mM NaCl, 1.5mM MgCl2, 5mM DTT) for 30 mins by rotating the mixture at room temperature. Agarose beads were washed three times with buffer B and incubated with non-biotinylated DNA (prey, 0.1 μM) and 0.4 μM purified mFoxP3ΔN protein.

After incubation for 30 mins with rotation, bead-bound DNA was recovered using proteinase K (New England Biolabs), purified by QIAquick Nucleotide Removal kit (QIAGEN) and analyzed on 10% Novex TBE Gels (Invitrogen). DNA was visualized by Sybr Gold staining.

#### Electrophoretic Mobility Shift Assay (EMSA)

DNA (0.2μM) was mixed with the indicated amount of FoxP3 and NFAT in buffer (20mM HEPES pH 7.5, 150mM NaCl, 1.5mM MgCl2 and 2mM DTT), incubated for 10 min at 4 °C and analyzed on 3-12% gradient Bis-Tris native gels (Life Technologies) at 4 °C. After staining with SYBR Gold stain (Life Technologies), Sybr Gold fluorescence was recorded using iBright FL1000 (Invitrogen) and analyzed with iBright Analysis Software.

#### Negative stain EM

FoxP3ΔN (0.4μM) was incubated with DNA (0.05μM) in buffer (20mM HEPES pH 7.5, 150mM NaCl, 1.5mM MgCl2 and 2mM DTT) at RT for 10 mins. The samples were diluted 10-fold with buffer, immediately adsorbed to freshly glow-discharged carbon-coated grids (Ted Pella) and stained with 0.75% uranyl formate as described before18.sImages were collected using a JEM- 1400 transmission electron microscope (JEOL) at 50,000X magnification.

#### NMR experiments

NMR experiments were measured on a 700 mM sample of 13C-15N murine FoxP1 RBR-FKH (aa 433-579) at 25 °C. The buffer was 20 mM MES pH=6.0, NaCl 150 mM, TCEP 2 mM, 2H2O 10% v/v. Spectra were collected on a Bruker spectrometer operating at 700 MHz 1H Larmor frequency, equipped with a triple-channel 1H, 13C, 15N cryogenically cooled probe and z-shielded gradients. All data were processed using NmrPipe 19 and analyzed using CCPNmr Analysis 20. Backbone 1H, 15N and 13C resonances were assigned using a standard set of triple resonance experiments: HSQC, HNCA, HN(CO)CA, HNCO, HN(CA)CO, HN(CA)CB, CBCA(CO)NH. 3D experiments were recorded using non-uniform sampling (NUS), where Poisson-Gap sampling was used to select 10% of the Nyquist grid 21 and the NUS data was reconstructed using the hmsIST protocol 22. 80% of expected resonances could be assigned, including 98% of RBR resonances. HN, N, CO, Cα and Cβ chemical shifts were used to derive dihedral angle values and secondary structure propensities using the software TALOS-N 23. Notably, two conformations of the RBR-C regions were assigned: one with high helical propensity and one with highly disordered character. 1H-15N HSQC spectra were compared at 700 and 50 mM and it was found that the disordered conformation does not exist significantly at lower concentrations. Chemical shift assignments have been deposited in the BMRB under accession number 5301.

#### FoxP3/FoxP1 PD-seq

Binding reactions were carried out by incubating 0.4 µM purified MBP-FoxP3ΔN or MBP-FoxP1ΔN protein with 0.1 µM dsDNA library in binding buffer (20 mM Tris-HCl pH 7.5, 100 mM NaCl, 1.5 mM MgCl₂) for 30 minutes at room temperature. DNA-protein complexes were captured using amylose resin (New England Biolabs), followed by three washes with binding buffer to remove non-specifically bound DNA. Elution of bound DNA was performed by using proteinase K (NEB) and DNA cleanup using the QIAquick Nucleotide Removal Kit (Qiagen). Eluted DNA was amplified using PCR with barcoded primers compatible with Illumina sequencing (primer sequences listed in Supplementary Table S1), libraries were pooled and subjected to deep sequencing.

#### Random-rcFKHM library PD-seq analysis

PD library and input samples were sequenced on an Illumina NovaSeq 6000 platform, generating paired-end 150 bp reads to a depth of 20 million reads each. Raw reads were merged with BBMerge2 (v 38.84), combining R1 and R2 reads into single reads at average merge rates of 98.67% (input), 99.47% (FoxP1 PD), and 99.16% (FoxP3 PD). Biopython3 (v1.85) was used to extract trimmed sequences matching the structure NNNNNNN–TCGA–GTAAACA (site 1–gap– rcFKHM). Trimmed sequences were combined across replicates and were organized into a Pandas4 (v 2.2.3) DataFrame, with each row corresponding to a unique 7-nucleotide sequence (excluding any sequences containing ’N’), yielding exactly 47 sequences. FoxP3/Input fold enrichment values, calculated from the combined replicates, were used to sort sequences for consistent ordering across all plots.

To identify enriched sequence patterns, we identified 7-nucleotide motifs containing two variable positions, denoted as ’N’. For each of the 47 selected sequences, we systematically generated all possible combinations of two variable positions, resulting in 21 degenerate “2N motifs” per sequence. Duplicate motifs were then removed across all sequences to avoid redundancy. For each resulting 2N motif, we enumerated all 16 possible nucleotide combinations at the two ’N’ positions (4 possibilities at each position). The counts for these 16 fully specified sequences were summed to calculate the total count for the 2N motif.

Enrichment ratios (FoxP3/Input) were computed for each 2N motif by comparing their aggregate counts in the FoxP3 and input datasets. Plots related to PD-seq motif counts and enrichment were visualized using Matplotlib7 (v 3.9.4).

#### Random-random library PD-seq analysis

FoxP3 PD and FoxP1 PD samples were sequenced on an Illumina NextSeq 2000 platform, generating single-end 150 bp reads to a depth of ∼280 million reads each. Reads with Phred quality score below 20 in the variable regions were discarded, resulting in ∼200 million reads each. Biopython3 (v1.85) was employed to extract trimmed sequences matching the structure: NNNNNNN-TCGA-NNNNNNN (site 1– gap–site 2) and were organized into a Pandas4 (v 2.2.3) DataFrame, with each row corresponding to a unique 18-nucleotide sequence, resulting in 104,367,238 unique FoxP1 sequences and 91,730,585 unique FoxP3 sequences. The DataFrame included columns for the normalized PD-seq counts (in RPM) of each sequence in the FoxP1 and FoxP3 samples.

Using the top 10 sequences from the random-rcFKHM PD-seq (Figure 1E) as a reference, we generated all pairwise motif combinations, resulting in 100 combinations (10X10). For each pair, we examined four orientations: Head-to-Head (H-H), Tail-to-Tail (T-T), Head-to-Tail (H-T) and Tail-to-Head (T-H). This resulted in 400 unique 18-nt long paired motifs, each with a fixed 4-nucleotide spacer “TCGA” between them.

Since the binding positions of FoxP3 in the random+random library were no longer fixed (unlike in the FKHM+random library), we scanned for motifs with -2, -1, 0, +1, and +2 nt offsets around the central gap. Each 18-nt sequence was transformed into five shifted variants that simulate 0, ±1, and ±2 base pair offsets around the central spacer (see Figure 2A). Each transformed sequence in FoxP3 and FoxP1 was checked against the precomputed dictionary containing all 400 paired motifs. If any shifted version matched a motif pair, the original read was counted as supporting that motif interaction. Since fixed sequences in the 4-nt gap and flanking regions introduced bias when motifs were counted during shifting, we computed the baseline likelihood of observing each motif pair under random offsets. We then normalized the observed motif frequency in PD samples by dividing by these baseline probabilities. Normalized frequencies were visualized as heatmaps and bar graphs using Matplotlib5 (v 3.9.4).

#### Motif analysis using in vivo dataset (ChIP-seq, CNR-seq, ATAC-seq, ChIP-exo-seq)

As in our previous work24, FoxP3 CNR-seq25,26 and FoxP3 Treg ChIP-seq16,27 datasets were trimmed by trimmomatic (v 0.3.6)12, aligned to mm10 using Bowtie27 (v2.3.4.3) and sorted by read position using SAMtools8 (v1.9). Peaks were called using MACS39 (v3.0.3) with a q-value cutoff of 0.001. Treg ATAC-seq peaks were downloaded from Immgen project28, and merged using Bedtools10 (v 2.31.0), which were referred to as open chromatin regions (OCRs). FoxP1 ChIP-seq28 peaks were called using the same pipeline, but with a q-value cutoff of 0.01, which were chosen to yield a peak count comparable to FoxP3. Finally, FoxP3 ChIP-exo peaks were defined as ±10 bp around reported ChIP-exo summits29. All peaks that overlap with blacklist regions30 were removed. This led to final sets of FoxP3 ChIP peaks (n = 25,724), CNR peaks (n = 18,552), ChIP-exo peaks (n = 4,307), FoxP1 ChIP peaks (n = 35,767) and ATAC peaks (n = 502,437).

To evaluate the presence of motif pairs in ChIP peaks, we extracted the genomic sequence from the GRCm38.p6 mouse reference genome31 for each peak and placed them in Pandas4 (v 2.2.3) DataFrames. Each sequence was scanned for exact matches to the 400 motifs identified in Figure 2C (100 motif pairs in H-H, H-T, T-H and T-T configurations). Note that only the positive strand of the genomic sequence was used to identify sequence matches. Matches within a given motif pattern were non-overlapping, as the search advanced past each match before continuing, to prevent inflated counts from repetitive or sliding matches of the same motif.

However, overlaps between different motif patterns were allowed to avoid search-order bias and ensure all valid matches were captured. To control for baseline motif frequency, we compared the frequency of each motif pair in FoxP3 ChIP peaks (or CNR or ChIP-exo peaks) to that in OCRs. We first normalized motif counts in ChIP peaks and OCRs with per-million base pair in their respective total peak lengths, and used exact binomial test32 to identify motifs that have significantly higher normalized counts in ChIP peaks than in OCRs (p-value < 0.05). For each of these motifs, the adjusted fold change (FC) of normalized counts in ChIP peaks over OCRs were visualized in the form of diameter scale in Figure 3A and heatmap in Figure 3B using Matplotlib5 (v 3.9.4).

Motif gap size analysis was performed using similar approaches, where normalized counts (per million bp of their respective total peak lengths) of motif pairs with a gap size of 1-10 nt were measured in ChIP peaks and OCRs, and were plotted using Matplotlib7 (v 3.9.4).

#### Analysis of relationships between TnG repeats and paired motifs

Defining genomic loci with TnG repeats FIMO from MEME suite (v5.5.8)13 was used to identify TnG-repeat-like elements. The TnG-repeat motif in our previous report14 was used as a query motif24, and a search was performed against the mouse (GRCm38) genome, FoxP3 ChIP peaks or OCRs. A p-value cutoff of 8e-5 was used in FIMO results, and FIMO regions with repetitive 6TGs, 6ACs, 6TAs, 6TCs, 6AGs, 6CGs, 12Ts, 12Gs, 12Cs or 12As were removed to be consistent with FoxP3’s sequence specificity14. The filtered TnG regions were exported using the python pyranges15 (v 0.1.4) package.

Relationships between TnG repeats and H-T/T-H motifs To examine the relationship between TnG repeats and H-T/T-H pairs of group 1-group 1 (G1-G1) motifs, we first compiled the genomic coordinates of H-T and T-H motif pairs within FoxP3 ChIP peaks using pyranges15 (v 0.1.4) package. Only the G1-G1 motifs that are significantly enriched (p<0.05) in FoxP3 ChIP peaks over OCRs were considered. We then calculated the percentage of H–T and T–H motifs that overlapped with TnG repeats defined above and plotted the values using Matplotlib5 (v 3.9.4).

#### Relationships between TnG/H–H and differential gene expression

RNA-seq datasets25,26,33 were acquired by mapping the reads to mm10 reference using splice-aware STAR mapper34, and exon reads counted using gencode mm10 annotation35. Deseq2 was used to calculate adjusted p-value and Log2FC values36. Genes were stratified into significant (adjusted p-value < 0.05) and non-significant groups. To quantify the enrichment of TnG or H–H features near these genes, we measured feature density (bp per Mb) within distance windows ranging from 1 kb to 1 Mb from each gene. Densities were computed using pyranges (v0.1.4).

Separately, to evaluate whether proximity to TnG or H–H features explained differential expression, the nearest distance from each gene to TnG repeats or H–H motifs was measured. Genes were grouped by adjusted p-value bins and log2 fold change direction, and the proportion of genes in each bin was calculated.

Relationships between TnG repeats and H-H motifs To assess how frequently H–H motifs and TnG repeats co-localize within the same FoxP3-bound regions or OCRs, each FoxP3 ChIP peaks or ATAC peaks examined for the presence of TnG repeats and H-H motifs using the pyranges15 (v 0.1.4) package.

To investigate the spatial and orientational relationships between H-H motifs and nearest TnG repeats, the genomic center and borders of each H-H motif were calculated, and the nearest TnG repeat was identified based on border-to- border distance. H-H motifs that had overlapping coordinates due to similar motif structure were merged to avoid duplicates. If the H-H motif coordinates overlapped with the coordinates of a TnG repeat, the border-to-border distance was recorded as zero. Otherwise, we measured the border-to-border distance as positive when the TnG occurred downstream (from the end of the H-H to the start of the TnG), and negative when it occurred upstream (from the end of the TnG to the start of the H-H). To assess the orientation of the nearest TnG repeat, we used strand information associated with the overlapping TnG region identified via the FIMO13 search. In cases where the TnG (positive strand) appeared upstream of the H-H, or CAn (negative strand) appeared downstream, the orientation was considered “aligned”. For comparison, the same analysis was performed in FoxP3 ChIP peaks and FoxP3-free OCRs (n=328,369). The latters were defined as Treg ATAC- seq37 peaks located at least 10 kb from any FoxP3 ChIP-seq peak and with FoxP3 ChIP-seq signal below the minimum signal in FoxP3 ChIP peaks, as measured by area under curve (AUC) for a 200 bp window around the center of each TnG repeat.

To compare FoxP3 binding to TnG repeats with and without adjacent H-H motifs, we calcualted FoxP3 ChIP-seq AUC for a 200 bp window around the center of each TnG repeat. TnG repeats with 0 nt gap with nearest H-H motif were compared against those located at least 1 kb away from any H–H motif.

#### Genomic feature and histone ChIP-seq analysis

Genomic feature analysis was performed using ChIPseeker38. To compare H3K4me3, H3K27ac and ATAC signal intensity, H3K4me3 and H3K27ac ChIP–seq and ATAC–seq data16 were trimmed by trimmomatic (v0.3.6)12 and mapped to the mm10 genome using Bowtie27 (v2.3.4.3). The intensity was calculated within 5 kb upstream and downstream of centers of TnG repeats and centers of H–H motifs using deeptools17 (v 3.5.6) bamCoverage and computeMatrix.

### Data and Code Availability

FoxP3 and FoxP1 pulldown-seq data have been deposited to the GEO database with the accession code of GSE294472. The NMR data has been deposited in the Biological Magnetic Resonance Bank (BMRB) with accession code of 5301. The custom codes used in this manuscript is deposited at Zenodo (DOI: 10.5281/zenodo.15306890). Any additional information required to reanalyze the data reported in this paper is available from the lead contact upon request.

---

**Figure S1.**
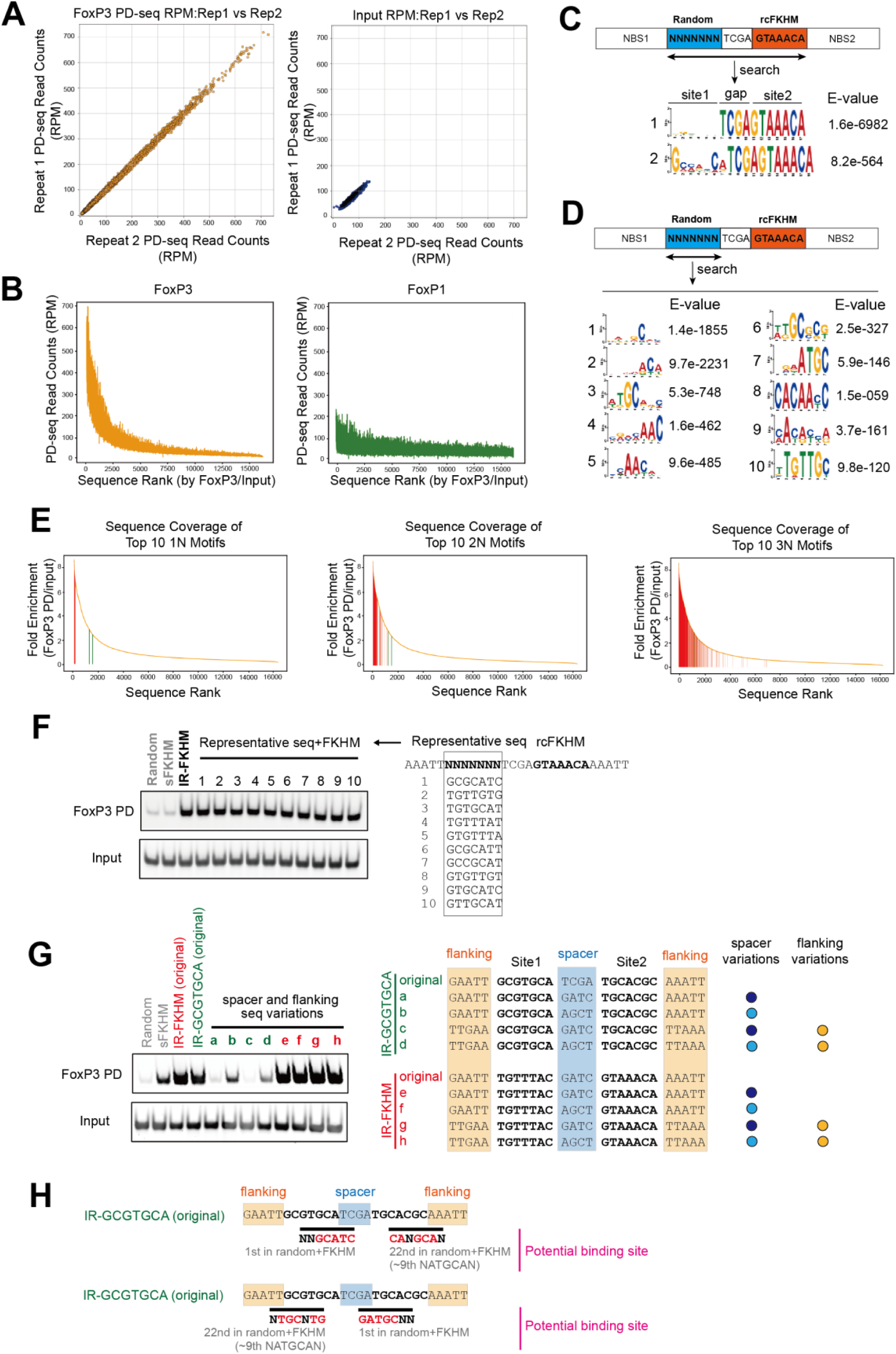
PD-seq analysis of FoxP3 and FoxP1 A. Scatter plots showing reproducibility between two biological replicates of FoxP3 PD-seq (left) and input control (right). Read counts (in RPM) for replicate 1 and replicate 2 are plotted on the x- and y-axes, respectively, demonstrating high concordance for FoxP3 PD-seq. B. PD-seq read count distributions for FoxP3 (left) and FoxP1 (right), with sequences ranked in descending order of FoxP3/Input enrichment (same rank order as in Figure 1B). FoxP3 RPM values show a steep decline, mirroring the enrichment distribution and confirming that highly enriched sequences are also abundant in the pull-downs ample. In contrast, FoxP1 RPM values are more uniform across the same sequence ranks, suggesting low selectivity for site1. C. De novo motif discovery using MEME to search sequences covering the 14 nt region encompassing site 1, the gap and site 2 (rcFKHM) failed to capture the variable sequences in site 1 due to overwhelming signal from the fixed sequence in the gap and site 2. D. De novo motif discovery using MEME on sequences containing only site 1. E. Rank-ordered enrichment plots (in descending order of FoxP3/Input enrichment) where sequences matching the top 10 motifs with degenerate bases at 1, 2, and 3 positions (1N, 2N, and 3N motifs) were indicated with red lines. Allowing 2N degeneracy achieves broad coverage of top-enriched sequences (top 10% of enrichment values) while maintaining resolution, which justifies its use in our motif analysis. Green lines mark sequences matching the putative GCGYGCH motif identified from the random–random library (Table S2), which was not enriched in the random–rcFKHM library. F. FoxP3 PD was performed using MBP-tagged FoxP3^ΔN^ (0.4 μM) to compare its binding affinity to DNA (0.1 μM) containing representative sequences of the top 10 2N-motifs, each paired with the reverse complement of FKHM (rcFKHM, GTAAACA). Numbers next to individual motifs indicate rank orders based on fold enrichment in Figure 1E. G. Spacer and flanking sequence requirements for IR-GCGTGCA vs. IR-FKHM binding. DNA constructs containing IR-GCGTGCA (green) or IR-FKHM (red) were tested in FoxP3 PD assays, performed as in (F). “Original” sequences correspond to motifs with spacer and flanking regions derived from the random-random library. Mutations in the spacer or flanking sequences abolished FoxP3 binding to IR-GCGTGCA, whereas IR-FKHM binding was largely unaffected. H. Schematic showing that IR-GCGTGCA within the random–random library harbors two inverted pairs of high-ranking motifs from Figure 1 at ±2 shifted positions. These overlapping motifs likely account for the apparent enrichment of IR-GCGYGCH.

**Figure S2.**
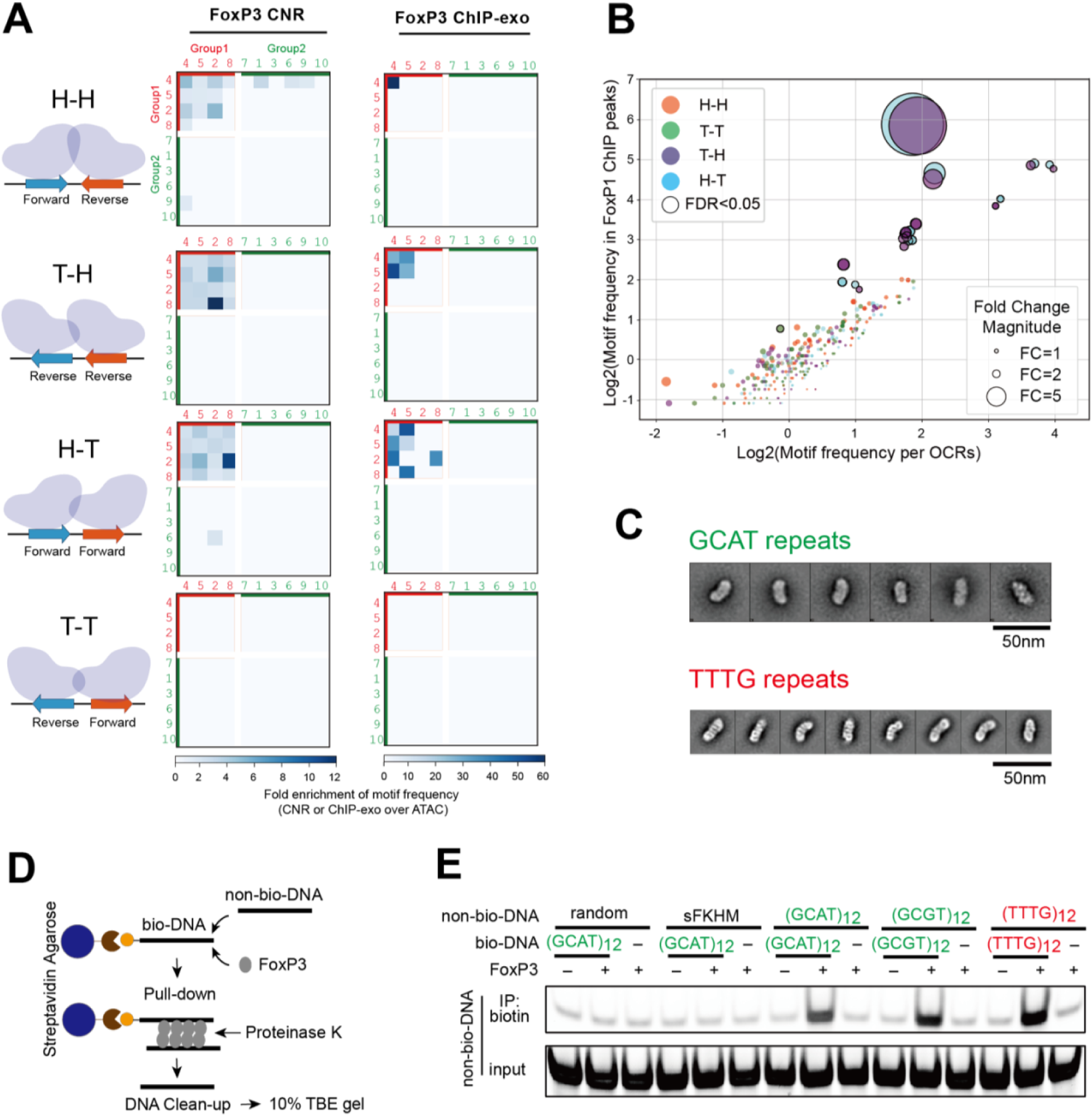
FoxP3 recognizes both H-H motifs and TnG repeats in Tregs. A. **Heatmaps showing fold change of the motif frequencies in FoxP3-bound sites over OCRs, grouped based on the motif pair orientation. The top 10 motifs are further categorized into group 1 (G1, red) and group 2 (G2, green) sequences. FoxP3** CNR-seq data are from Tregs (left) and ChIP-exo from mESCs (right). Only the motif pairs that are significantly enriched in FoxP3-bound sites (p<0.05, exact binomial test) are shaded. B. Log2-transformed motif frequencies in FoxP1 ChIP-seq peaks (35,767 peaks) vs. OCRs (based on ATAC-seq, 502,437 peaks). Each point represents one of the 400 motif pairs (10 motifs x 10 motifs x 4 orientation combinations) with a 4-nt gap, colored by the motif orientation relative to the reference genome (positive strand): H-H, T-H, H-T, and T-T. Motif frequency was defined as the number of motif occurrence within all ChIP-peaks (or ATAC-peaks) divided by the total length (in million bp) of those peaks. Dot size reflects fold change of motif frequency in ChIP-peaks over OCRs. Motif pairs significantly enriched in FoxP3-bound regions (p<0.05, exact binomial test) are outlined in black and are further analyzed in Figure 3B. C. Representative 2D class averages from negative-stain electron microscopy of FoxP3^ΔN^ in complex with (GCAT)_15_ (top) and (TTTG)_18_ (bottom). FoxP3 multimers form on both TTTG and CGAT repeats with similar morphologies. Consistent with DNA length, FoxP3 multimers on both sequences measure ∼25 nm. D. Schematic of DNA bridging assay. Biotinylated DNA (bio-DNA, 82 bp), pre-conjugated to streptavidin agarose beads, and non-biotinylated DNA (non-bio-DNA, 48 bp) were mixed at a 1:1 ratio (0.1 μM each), incubated with FoxP3^ΔN^ (0.4 μM) and subjected to pull-down. Co- purified non-bio-DNA was then recovered by proteinase K digestion and analyzed on TBE gels (see Methods for details). E. DNA bridging assay using indicated bio-DNA and non-bio-DNA sequences.

**Figure S3.**
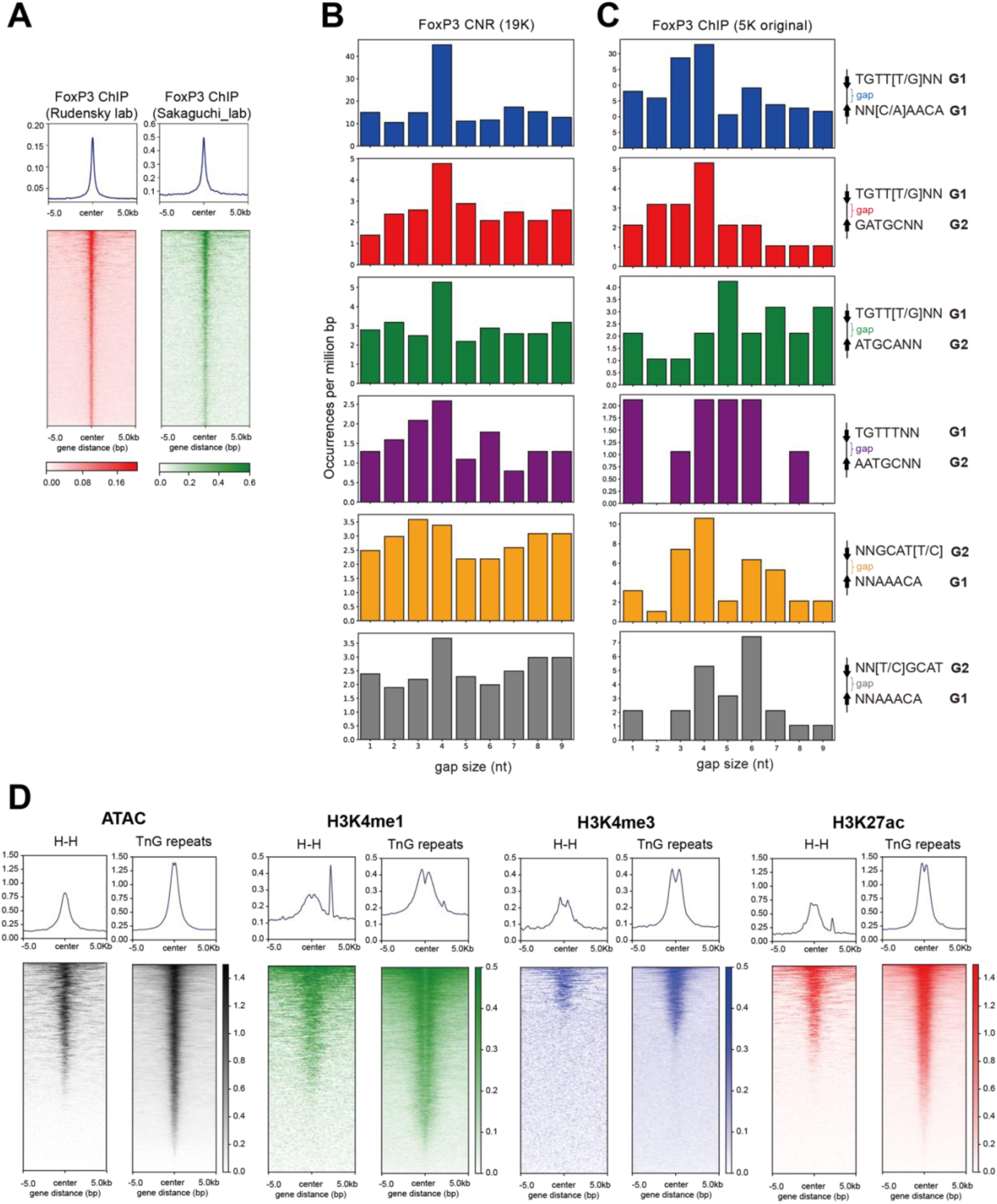
Analysis of FoxP3-bound H-H motifs in Tregs. A. FoxP3 ChIP-seq signal across ±5 kb centered on H-H motif-containing loci in Tregs from Rudensky lab (PMID:23021222) and Sakaguchi lab (PMID: 29044238). Each row represents a genomic region, ranked using the same order in both heatmaps. Average signal profile is shown above each heatmap. B. Gap size distributions for the indicated motif pairs across **FoxP3 CnR-seq peaks (n=18,552)**. C. Gap size distributions for the indicated motif pairs across the original **FoxP3 ChIP-seq peaks (n=5,047) used in PMID:** 35926508. D. Signal intensities for ATAC-seq and histone ChIP-seq (H3K27ac, H3K4me1, and H3K4me3) across ±5 kb centered on FoxP3-bound TnG repeats (n=19,145) and H–H motifs (n=2,375). Each row represents a genomic region. Average signal profile is shown above each heatmap.

**Figure S4.**
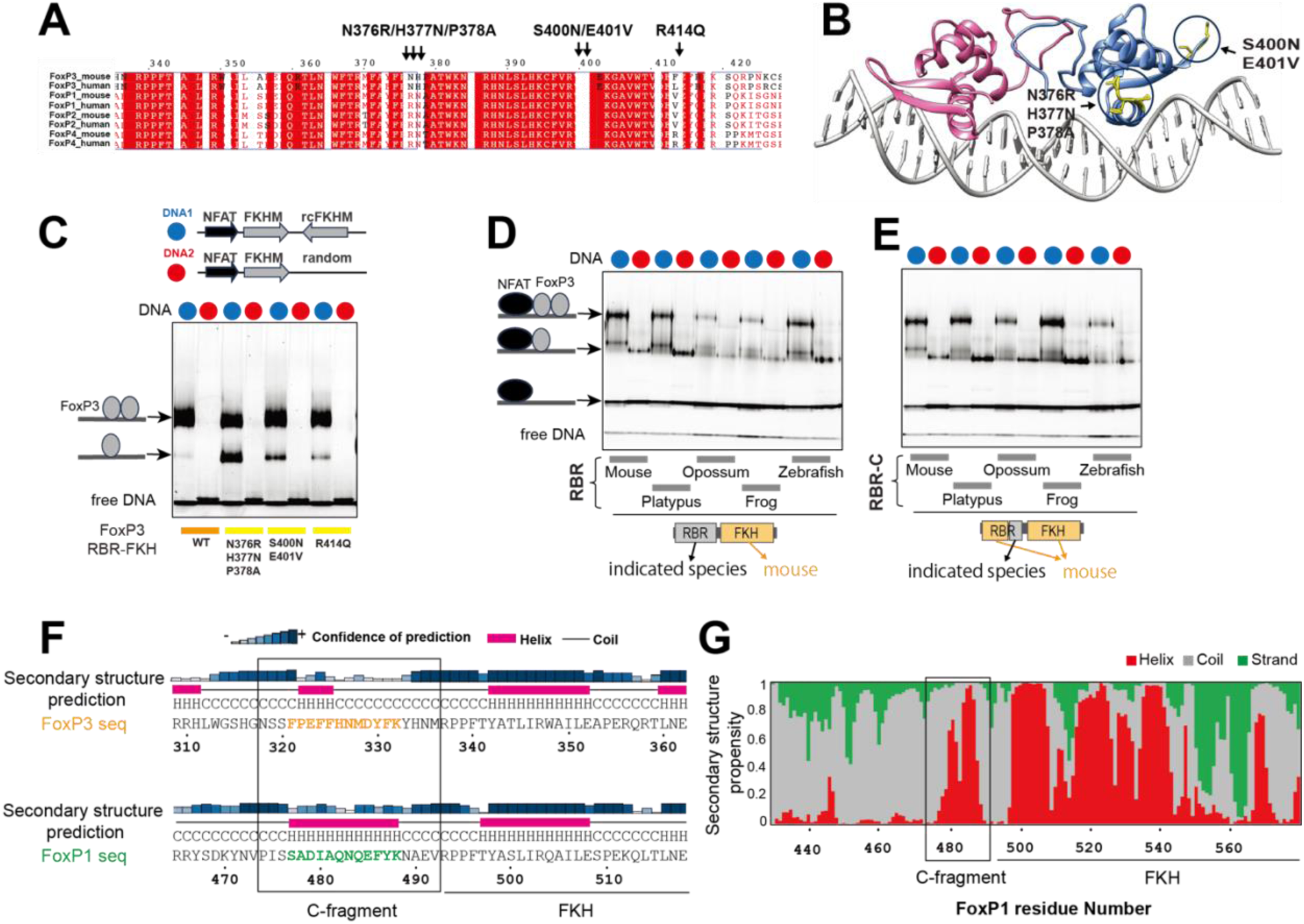
RBR-C is the key determinant for H-H dimerization of FoxP3 A. Sequence alignment of the FKH domain of mouse and human FoxP TFs, with moderately conserved residues indicated in red font and highly conserved residues highlighted in red. Arrows indicate divergent residues between FoxP3 and FoxP1/2/4. B. Structure highlighting the divergent residues in the FKH domain indicated in (A). C. Native gel shift assay of WT and mutant FoxP3^RBR-FKH^ (0.8 μM, NusA-tagged) with IR-FKHM (blue) or sFKHM (red) DNA (0.2 μM). Mutations in the divergent residues in (A) had minimal impact on FoxP3’s preference for IR-FKHM over sFKHM, although they commonly reduced FoxP3’s dimerization on IR-FKHM to varying degrees. D-E. Native gel shift assays using various chimeric RBR-FKH constructs, in which the RBR or RBR- C domain of mouse FoxP3 was replaced with that from different FoxP3 orthologs. FoxP3 (0.8 μM) was incubated with IR-FKHM (blue) or sFKHM (red) DNA (0.2 μM) in the presence of NFAT (0.4 μM). Although swapping the FoxP3 RBR loop with those from other species altered the overall DNA-binding affinity, it did not affect the preferred binding mode; all constructs bound IR-FKHM as a dimer and sFKHM as a monomer, mirroring the behavior of unswapped mouse FoxP3. F. Secondary structure prediction (by PSIPRED) indicated an α-helix (purple) in the RBR-C region of FoxP1 (bottom, highlight in green), whereas FoxP3 RBR-C (top, highlight in yellow) was predicted to contain only a short, four–amino acid helical stretch. G. NMR experiments indicated a strong α-helical propensity (red) for FoxP1 residues 477–488 within RBR-C fragment, consistent with the predicted secondary structure.

## REFERENCES

1 Lambert, S. A. et al. The Human Transcription Factors. Cell 172, 650–665 (2018). 10.1016/j.cell.2018.01.029

2 Nitta, K. R. et al. Conservation of transcription factor binding specificities across 600 million years of bilateria evolution. Elife 4 (2015). 10.7554/eLife.04837

3 Leng, F. et al. The transcription factor FoxP3 can fold into two dimerization states with divergent implications for regulatory T cell function and immune homeostasis. Immunity 55, 1354–1369 e1358 (2022). 10.1016/j.immuni.2022.07.002

4 Zhang, W. et al. FOXP3 recognizes microsatellites and bridges DNA through multimerization. Nature 624, 433–441 (2023). 10.1038/s41586-023-06793-z

5 Leng, F. et al. Ultrastable and versatile multimeric ensembles of FoxP3 on microsatellites. Mol Cell (2025). 10.1016/j.molcel.2025.03.005

6 Dai, S., Qu, L., Li, J. & Chen, Y. Toward a mechanistic understanding of DNA binding by forkhead transcription factors and its perturbation by pathogenic mutations. Nucleic Acids Res 49, 10235–10249 (2021). 10.1093/nar/gkab807

7 Wu, Y. et al. FOXP3 controls regulatory T cell function through cooperation with NFAT. Cell 126, 375–387 (2006). 10.1016/j.cell.2006.05.042

8 Bandukwala, H. S. et al. Structure of a domain-swapped FOXP3 dimer on DNA and its function in regulatory T cells. Immunity 34, 479–491 (2011). 10.1016/j.immuni.2011.02.017

9 Arora, S. et al. Joint sequence & chromatin neural networks characterize the differential abilities of Forkhead transcription factors to engage inaccessible chromatin. bioRxiv (2023). 10.1101/2023.10.06.561228

10 Ono, M. et al. Foxp3 controls regulatory T-cell function by interacting with AML1/Runx1. Nature 446, 685–689 (2007). 10.1038/nature05673

11 Ramirez, R. N., Chowdhary, K., Leon, J., Mathis, D. & Benoist, C. FoxP3 associates with enhancer-promoter loops to regulate T(reg)-specific gene expression. Sci Immunol 7, eabj9836 (2022). 10.1126/sciimmunol.abj9836

12 Liu, Z., Lee, D. S., Liang, Y., Zheng, Y. & Dixon, J. R. Foxp3 orchestrates reorganization of chromatin architecture to establish regulatory T cell identity. Nat Commun 14, 6943 (2023). 10.1038/s41467-023-42647-y

13 Song, X. et al. Structural and biological features of FOXP3 dimerization relevant to regulatory T cell function. Cell Rep 1, 665–675 (2012). 10.1016/j.celrep.2012.04.012

14 van der Veeken, J. et al. The Transcription Factor Foxp3 Shapes Regulatory T Cell Identity by Tuning the Activity of trans-Acting Intermediaries. Immunity 53, 971–984 e975 (2020). 10.1016/j.immuni.2020.10.010

15 Kitagawa, Y. et al. Guidance of regulatory T cell development by Satb1-dependent super-enhancer establishment. Nat Immunol 18, 173–183 (2017). 10.1038/ni.3646

16 de Boer, C. G. & Taipale, J. Hold out the genome: a roadmap to solving the cis-regulatory code. Nature 625, 41–50 (2024). 10.1038/s41586-023-06661-w

17 Bailey, T. L., Johnson, J., Grant, C. E. & Noble, W. S. The MEME Suite. Nucleic Acids Res 43, W39–49 (2015). 10.1093/nar/gkv416

18 Nakagawa, S., Gisselbrecht, S. S., Rogers, J. M., Hartl, D. L. & Bulyk, M. L. DNA-binding specificity changes in the evolution of forkhead transcription factors. Proc Natl Acad Sci U S A 110, 12349–12354 (2013). 10.1073/pnas.1310430110

19 Heng, T. S., Painter, M. W. & Immunological Genome Project, C. The Immunological Genome Project: networks of gene expression in immune cells. Nat Immunol 9, 1091– 1094 (2008). 10.1038/ni1008-1091

20 Zhu, Z. et al. FOXP1 and KLF2 reciprocally regulate checkpoints of stem-like to effector transition in CAR T cells. Nat Immunol 25, 117–128 (2024). 10.1038/s41590-023-01685-w

21 Jager, C. et al. Inducible protein degradation reveals inflammation-dependent function of the T(reg) cell lineage-defining transcription factor Foxp3. Sci Immunol 10, eadr7057 (2025). 10.1126/sciimmunol.adr7057

22 Zemmour, D. et al. Single-cell analysis of FOXP3 deficiencies in humans and mice unmasks intrinsic and extrinsic CD4(+) T cell perturbations. Nat Immunol 22, 607–619 (2021). 10.1038/s41590-021-00910-8

## REFERENCES

1 Tang, G. et al. EMAN2: an extensible image processing suite for electron microscopy. J Struct Biol 157, 38–46 (2007). 10.1016/j.jsb.2006.05.009

2 Bushnell, B., Rood, J. & Singer, E. BBMerge - Accurate paired shotgun read merging via overlap. PLoS One 12, e0185056 (2017). 10.1371/journal.pone.0185056

3 Cock, P. J. et al. Biopython: freely available Python tools for computational molecular biology and bioinformatics. Bioinformatics 25, 1422–1423 (2009). 10.1093/bioinformatics/btp163

4 McKinney, W. in Proceedings of the 9th Python in Science Conference (ed S.; Millman van der Walt, J.) 51–56 (2010).

5 Hunter, J. Matplotlib: A 2D graphics environment. Computing in Science & Engineering 9 90–95 (2007). 10.1109/MCSE.2007.55

6 Harris, C. R. et al. Array programming with NumPy. Nature 585, 357–362 (2020). 10.1038/s41586-020-2649-2

7 Langmead, B. & Salzberg, S. L. Fast gapped-read alignment with Bowtie 2. Nat Methods 9, 357–359 (2012). 10.1038/nmeth.1923

8 Danecek, P. et al. Twelve years of SAMtools and BCFtools. Gigascience 10 (2021). 10.1093/gigascience/giab008

9 Zhang, Y. et al. Model-based analysis of ChIP-Seq (MACS). Genome Biol 9, R137 (2008). 10.1186/gb-2008-9-9-r137

10 Quinlan, A. R. & Hall, I. M. BEDTools: a flexible suite of utilities for comparing genomic features. Bioinformatics 26, 841–842 (2010). 10.1093/bioinformatics/btq033

11 Leinonen, R., Sugawara, H., Shumway, M. & International Nucleotide Sequence Database, C. The sequence read archive. Nucleic Acids Res 39, D19–21 (2011). 10.1093/nar/gkq1019

12 Bolger, A. M., Lohse, M. & Usadel, B. Trimmomatic: a flexible trimmer for Illumina sequence data. Bioinformatics 30, 2114–2120 (2014). 10.1093/bioinformatics/btu170

13 Grant, C. E., Bailey, T. L. & Noble, W. S. FIMO: scanning for occurrences of a given motif. Bioinformatics 27, 1017–1018 (2011). 10.1093/bioinformatics/btr064

14 Bailey, T. L., Johnson, J., Grant, C. E. & Noble, W. S. The MEME Suite. Nucleic Acids Res 43, W39-49 (2015). 10.1093/nar/gkv416

15 Stovner, E. B. & Saetrom, P. PyRanges: efficient comparison of genomic intervals in Python. Bioinformatics 36, 918–919 (2020). 10.1093/bioinformatics/btz615

16 Kitagawa, Y. et al. Guidance of regulatory T cell development by Satb1-dependent super-enhancer establishment. Nat Immunol 18, 173–183 (2017). 10.1038/ni.3646

17 Ramirez, F. et al. deepTools2: a next generation web server for deep-sequencing data analysis. Nucleic Acids Res 44, W160–165 (2016). 10.1093/nar/gkw257

18 Ohi, M., Li, Y., Cheng, Y. & Walz, T. Negative Staining and Image Classification - Powerful Tools in Modern Electron Microscopy. Biol Proced Online 6, 23–34 (2004). 10.1251/bpo70

19 Delaglio, F. et al. NMRPipe: a multidimensional spectral processing system based on UNIX pipes. J Biomol NMR 6, 277–293 (1995). 10.1007/bf00197809

20 Vranken, W. F. et al. The CCPN data model for NMR spectroscopy: development of a software pipeline. Proteins 59, 687–696 (2005). 10.1002/prot.20449

21 Hyberts, S. G., Takeuchi, K. & Wagner, G. Poisson-gap sampling and forward maximum entropy reconstruction for enhancing the resolution and sensitivity of protein NMR data. J Am Chem Soc 132, 2145–2147 (2010). 10.1021/ja908004w

22 Hyberts, S. G., Milbradt, A. G., Wagner, A. B., Arthanari, H. & Wagner, G. Application of iterative soft thresholding for fast reconstruction of NMR data non-uniformly sampled with multidimensional Poisson Gap scheduling. J Biomol NMR 52, 315-327 (2012). 10.1007/s10858-012-9611-z

23 Shen, Y. & Bax, A. Protein backbone and sidechain torsion angles predicted from NMR chemical shifts using artificial neural networks. J Biomol NMR 56, 227–241 (2013). 10.1007/s10858-013-9741-y

24 Zhang, W. et al. FOXP3 recognizes microsatellites and bridges DNA through multimerization. Nature 624, 433–441 (2023). 10.1038/s41586-023-06793-z

25 van der Veeken, J. et al. The Transcription Factor Foxp3 Shapes Regulatory T Cell Identity by Tuning the Activity of trans-Acting Intermediaries. Immunity 53, 971–984 e975 (2020). 10.1016/j.immuni.2020.10.010

26 Liu, Z., Lee, D. S., Liang, Y., Zheng, Y. & Dixon, J. R. Foxp3 orchestrates reorganization of chromatin architecture to establish regulatory T cell identity. Nat Commun 14, 6943 (2023). 10.1038/s41467-023-42647-y

27 Samstein, R. M. et al. Foxp3 exploits a pre-existent enhancer landscape for regulatory T cell lineage specification. Cell 151, 153–166 (2012). 10.1016/j.cell.2012.06.053

28 Zhu, Z. et al. FOXP1 and KLF2 reciprocally regulate checkpoints of stem-like to effector transition in CAR T cells. Nat Immunol 25, 117–128 (2024). 10.1038/s41590-023-01685-w

29 Yamada, N., Rossi, M. J., Farrell, N., Pugh, B. F. & Mahony, S. Alignment and quantification of ChIP-exo crosslinking patterns reveal the spatial organization of protein-DNA complexes. Nucleic Acids Res 48, 11215–11226 (2020). 10.1093/nar/gkaa618

30 Amemiya, H. M., Kundaje, A. & Boyle, A. P. The ENCODE Blacklist: Identification of Problematic Regions of the Genome. Sci Rep 9, 9354 (2019). 10.1038/s41598-019-45839-z

31 Yates, A. D. et al. Ensembl 2020. Nucleic Acids Res 48, D682–D688 (2020). 10.1093/nar/gkz966

32 Virtanen, P. et al. SciPy 1.0: fundamental algorithms for scientific computing in Python. Nat Methods 17, 261–272 (2020). 10.1038/s41592-019-0686-2

33 Jager, C. et al. Inducible protein degradation reveals inflammation-dependent function of the T(reg) cell lineage-defining transcription factor Foxp3. Sci Immunol 10, eadr7057 (2025). 10.1126/sciimmunol.adr7057

34 Dobin, A. et al. STAR: ultrafast universal RNA-seq aligner. Bioinformatics 29, 15–21 (2013). 10.1093/bioinformatics/bts635

35 Frankish, A. et al. GENCODE reference annotation for the human and mouse genomes. Nucleic Acids Res 47, D766–D773 (2019). 10.1093/nar/gky955

36 Love, M. I., Huber, W. & Anders, S. Moderated estimation of fold change and dispersion for RNA-seq data with DESeq2. Genome Biol 15, 550 (2014). 10.1186/s13059-014-0550-8

37 Heng, T. S., Painter, M. W. & Immunological Genome Project, C. The Immunological Genome Project: networks of gene expression in immune cells. Nat Immunol 9, 1091–1094 (2008). 10.1038/ni1008-1091

38 Yu, G., Wang, L. G. & He, Q. Y. ChIPseeker: an R/Bioconductor package for ChIP peak annotation, comparison and visualization. Bioinformatics 31, 2382–2383 (2015). 10.1093/bioinformatics/btv145

